# Smoothie: Efficient Inference of Spatial Co-expression Networks from Denoised Spatial Transcriptomics Data

**DOI:** 10.1101/2025.02.26.640406

**Authors:** Chase Holdener, Iwijn De Vlaminck

## Abstract

Finding correlations in spatial gene expression is fundamental in spatial transcriptomics, as co-expressed genes within a tissue are linked by regulation, function, pathway, or cell type. Yet, sparsity and noise in spatial transcriptomics data pose significant analytical challenges. Here, we introduce Smoothie, a method that denoises spatial transcriptomics data with Gaussian smoothing and constructs and integrates genome-wide co-expression networks. Utilizing implicit and explicit parallelization, Smoothie scales to datasets exceeding 100 million spatially resolved spots with fast run times and low memory usage. We demonstrate how co-expression networks measured by Smoothie enable precise gene module detection, functional annotation of uncharacterized genes, linkage of gene expression to genome architecture, and multi-sample comparisons to assess stable or dynamic gene expression patterns across tissues, conditions, and time points. Overall, Smoothie provides a scalable and versatile framework for extracting deep biological insights from high-resolution spatial transcriptomics data.

## INTRODUCTION

Recent advances in high-resolution spatial transcriptomics enable profiling of millions of spatially resolved transcriptomes within a single tissue sample. Yet, analyzing these data remains challenging due to noise, data sparsity, and the fact that each “spot” can capture partial gene expression from multiple cells. A powerful but underutilized strategy for interpreting complex spatial RNA data is to measure weighted spatial gene co-expression networks, which capture how gene pairs co-express across physical locations in the tissue. Weighted co-expression network analysis was first introduced on microarray bulk RNA data^1,2^ and has been adapted for single cell RNA sequencing and low-resolution spatial RNA sequencing^3,4^. In spatial transcriptomics, analyzing spatial gene modules and their networks can identify novel gene associations, even among previously uncharacterized genes, discovering new gene functions and new gene annotations. Further, the gene correlation network provides a straightforward way to summarize the co-expression patterns of any gene set of interest, such as a gene family, a gene pathway, cell communication genes, transcription factor genes, or more. As we show here, gene correlation networks furthermore provide an avenue for harmonizing and cross-analyzing data from different tissues, a task for which few substantive solutions exist. Critically, constructing gene correlation networks does not require prior knowledge of tissue biology or architecture, and does not require companion data (e.g., imaging or single-cell transcriptomic data) or specialized preprocessing such as cell segmentation, which can be difficult to perform accurately^5^.

Given the importance of spatial gene correlation networks, it is essential to measure them with high accuracy and computational efficiency. Here, we introduce and validate *Smoothie*, an algorithm for fast and precise measurement of spatial gene correlation networks in high resolution spatial transcriptomics. Smoothie performs Gaussian smoothing on the data to address noise and sparsity, constructs genome-wide gene correlation networks, and then finds gene modules with graph-based clustering for downstream analyses. Smoothie has high computational efficiency, processing datasets exceeding 100 million spatial spots and over 20,000 genes—resulting in 200 million pairwise gene correlation calculations—all in about 1 hour on a standard high-performance CPU. Thus, Smoothie can handle large datasets from the newest spatial platforms.

We demonstrate how Smoothie accurately identifies modules of matching gene expression patterns in both the adult mouse cerebellum and the developing mouse embryo, where gene modules correspond to tissue regions, known cell types, and local genome positioning. We further use Smoothie to associate over 50 previously uncharacterized or predicted mouse genes with known cell types and spatial gene programs, and we spatially characterize the Slc transporter superfamily in the mouse embryo. Finally, we show how second-order gene correlations can be used to detect differences in erythroid and epidermis expression patterns across timepoints in late mouse embryonic development (E13.5-E16.5). Further, Smoothie makes the discovery that 15 members of the Fbxw gene family are uniquely expressed in the cumulus oocyte cluster cells in the ovary throughout mouse ovulation. In conclusion, Smoothie offers an efficient and versatile approach for biological discovery in spatial transcriptomics data by leveraging spatial gene correlation networks.

## RESULTS

### Smoothie precisely measures spatial gene correlations

Smoothie implements a three-step algorithm to output spatial gene pattern modules in an interpretable gene network format (**Fig. 1A**). The input is a spatial gene count matrix *X* with *N* spatial spots and *G* genes. In the first step, Smoothie applies Gaussian smoothing to remove fine-scale noise and to address data sparsity. Specifically, Smoothie moves a Gaussian distribution with standard deviation σacross the (*x, y*) locations in the dataset and replaces each gene expression value at each (*x, y*) location with a Gaussian weighted average of the nearby gene values (**Methods**). In the second step, Smoothie creates a pairwise gene correlation matrix *R* of size *G×G* by computing the Pearson correlation coefficient (PCC) between all pairs of genes in parallel. Finally, Smoothie constructs a weighted spatial gene correlation network *W*, with nodes being the set of genes *V*= {*g*_1_, *g*_2_,... *g*_*G*_} and edges being the set *E* = {(*g*_*i*_, *g*_*j*_, *R*_*ij*_) |*R*_*ij*_ >*PCC*\*}, where *PCC*\* is a cutoff for correlation significance determined by considering a spatially shuffled version of *X*(**Methods**). The network positions genes based on their pattern similarities with each other and enables grouping of genes into modules using graph clustering. Smoothie supports multiple downstream analyses, including identifying precise spatial gene modules, investigating individual genes through spatial correlations, connecting gene correlations with genome position, examining specific gene sets or families, and more (**Fig. 1B, Figure S2**). Additionally, Smoothie enables spatial gene module integration across datasets with mismatched coordinates and provides a way to rank genes as stable or variable spatial expression patterns across diverse conditions (**Fig. 1C**).

**Figure 1.**
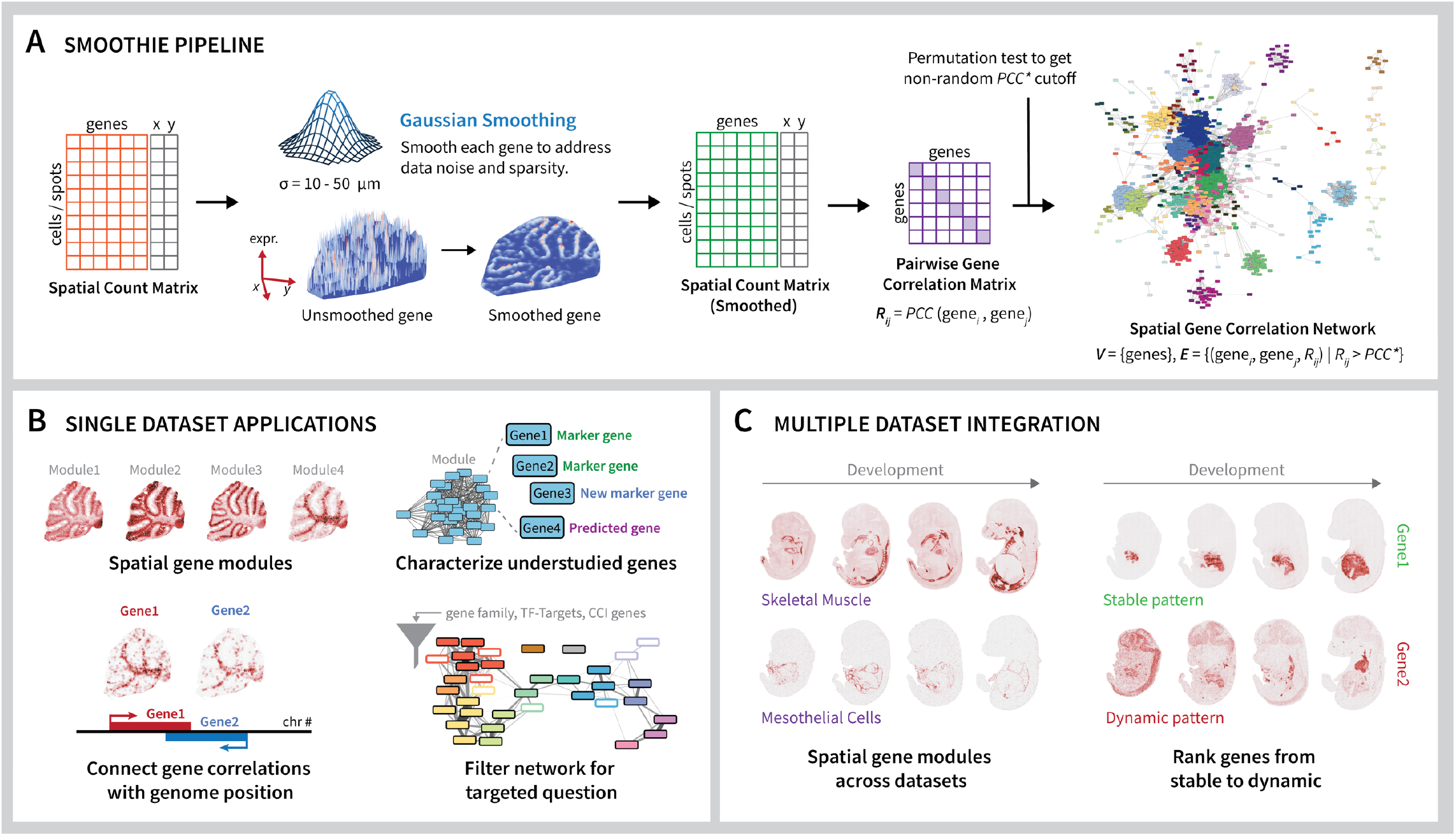
Overview of the Smoothie pipeline and its applications in one or many spatial datasets. **A**.Smoothie takes a spatial count matrix as input and performs efficient Gaussian smoothing on each gene using a 2D Gaussian with standard deviation ***σ***. Smoothie calculates the correlation of every smoothed gene pair to create a gene correlation matrix *R*. The gene correlation network with genes as vertices and weighted edges reflecting spatial gene correlations comes from the matrix *R*, using a spatially permuted count matrix to find a significant nonrandom correlation cutoff *PCC*\*. Graph clustering algorithms then identify network gene modules (**Methods**). **B**. Several powerful biological use cases follow from the gene correlation network. First, the network finds spatial gene modules grouped by pattern similarity. Second, gene modules characterize understudied or predicted genes by connecting them with known gene pathways, cell types, or tissue regions. Third, spatial gene correlations may reflect local genome positioning. Fourth, the gene correlation network may be filtered for any gene set of interest to investigate pattern diversity of the gene set members, such as a gene family, TFs and target genes, or cell-cell interaction (CCI) genes. **C**. Smoothie provides integration analyses for multiple spatial dataset experiments, such as finding spatial gene modules across datasets and ranking genes as stable or dynamic across datasets.

### Smoothie identifies spatial gene modules in the adult mouse cerebellum

We first applied Smoothie to the Slide-seq Puck_180819_12 cerebellum dataset^6^, which spans 32,701 10 μm-resolution spots (≥5 UMIs each) and 6,942 genes (≥50 detected spots). After applying Gaussian smoothing (30 μm Gaussian standard deviation), Smoothie detected 17,208 gene pairs with PCC > 0.35 (**Data S1, Methods**). We treated gene pairs as weighted edges in a correlation network and used weighted random network walks clustering^7^ to identify 10 modules of correlated genes (**Data S2**). Tightening the cutoff to PCC > 0.6 revealed 270 edges among 118 genes, and highlighted substructure in the correlation network (**Fig. 2A**). For example, Purkinje cell marker genes *Calb1, Pvalb, Car8*, and *Pcp4* (module 3) grouped in a closely connected cluster, as do oligodendrocyte markers *Trf, Mobp, Apod*, and *Mal* (module 4, **Fig. 2B**). The network also captured subtle intra-module differences of genes—within module 1, *Meg3* correlated more strongly to *Snap25*, whereas *Nap1l5* is more similar to *Cartpt* (**Fig. 2C**). Further, genes that bridge two modules together, such as *Itm2b* or *Sparcl1*, are evident and retrievable from the network. The 10 Smoothie modules from the PCC > 0.35 network reveal distinct gene patterns in the mouse cerebellum (**Fig. 2D**), many of which correspond to a specific cell subclass or subclasses identified in the cerebellum region of the Allen Brain Cell Atlas^8^ (**Fig. S5**). Modules 2, 3, 4, and 5 correspond to granule cells, Purkinje cells, oligodendrocytes, and choroid plexus cells, respectively. Module 6 is expressed primarily in neurons outside the cerebellum; module 7 is widely expressed across all cell types; and module 8 contains markers of non-neuronal cell types, particularly Bergmann glial cells, astrocytes, and ependymal cells. Modules 9 and 10 are smaller, containing only 3 and 2 genes respectively; module 9 appears to align with glutamatergic neurons and module 10 with monocytes.

**Figure 2.**
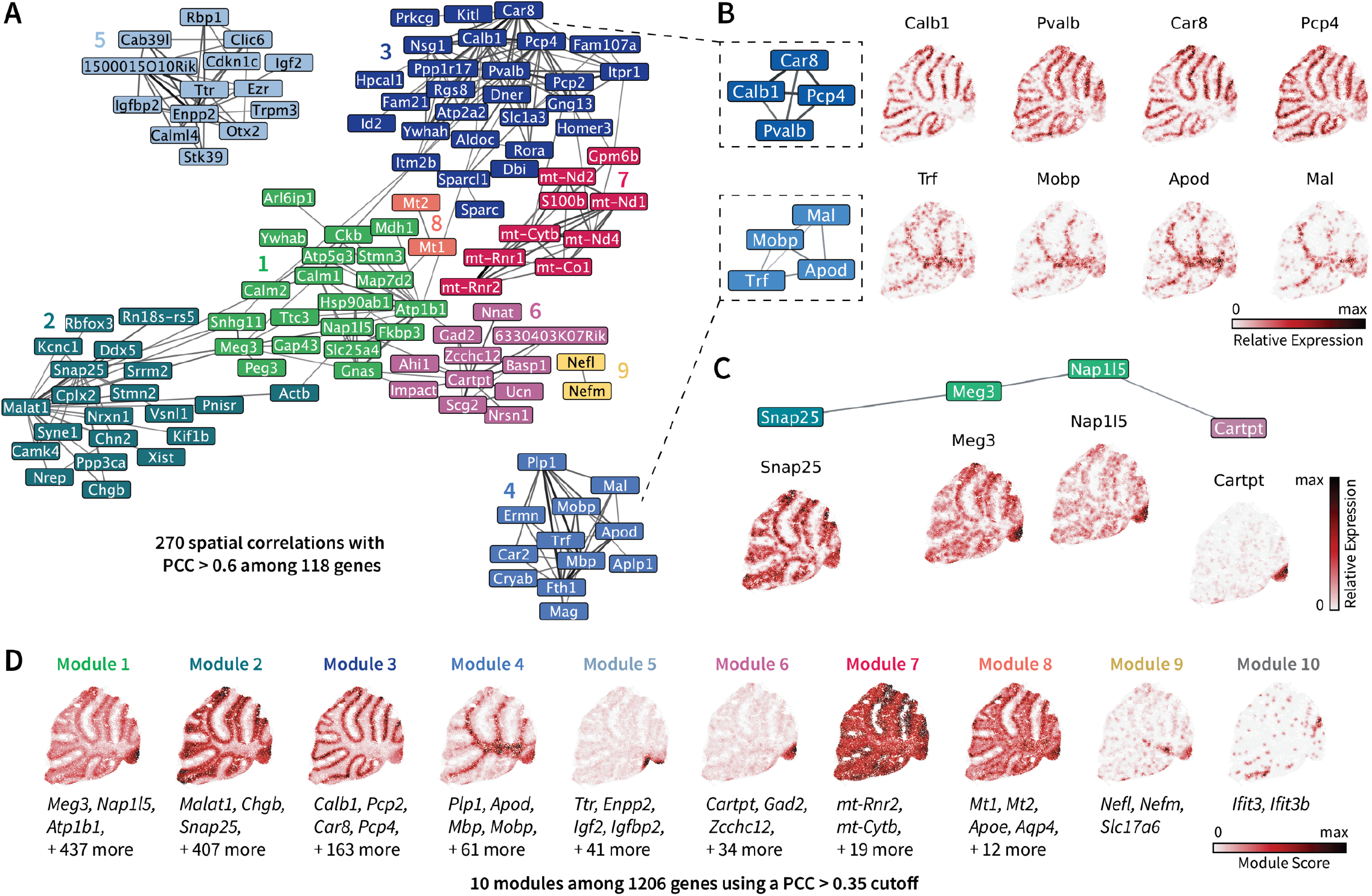
Smoothie identifies spatial gene modules in the adult mouse cerebellum. **A**.The spatial gene correlation network, representing genes as nodes and spatial gene correlations as edges. Edges are weighted by the Pearson correlation coefficient (PCC) between the two gene surfaces, and the width and darkness of the line reflects edge weight. Nodes are colored by their assigned module. Network cutoff PCC > 0.6. **B**. Example spatial expression patterns from tightly connected genes in the network. Plots show relative gene expression after normalization and smoothing. **C**. Example spatial expression patterns from connected genes spanning multiple modules. **D**. Module score plots for each of the 10 modules identified by Smoothie with network cutoff PCC > 0.35. The module score reflects average smoothed expression of all genes in the module after rescaling each gene’s maximum value to 1.

We next used these data to compare Smoothie to Hotspot^9^, a leading method for identifying spatial gene correlations. Hotspot detected 6 gene modules in the cerebellum dataset, that align with Smoothie’s modules 1–6 but did not recover modules 7–10 (**Fig. S5**). To examine pairwise gene similarity ranks, we focused on 3 genes (*Car8, Malat1*, and *Ttr*) and compared the top 300 most similar genes identified by both Smoothie and Hotspot (**Fig. S4**). While most top-ranked genes were shared by both methods, Smoothie identified several sparse genes (<500 UMIs) whose expression patterns visibly correlated with each gene of interest, but which Hotspot disregarded due to low autocorrelation scores. Conversely, Hotspot sometimes incorrectly prioritized genes whose expression diverged from the gene of interest. Finally, runtime tests varying the number of genes and spatial locations confirmed Smoothie was significantly faster than Hotspot across all tested inputs, with over 100-fold runtime improvements for the larger tested inputs (**Fig. S3**).

### Smoothie finds new gene modules and new gene annotations in the E16.5 mouse embryo

Next, we applied Smoothie to the larger and more complex 0.5 μm-resolution dataset of a E16.5 mouse embryo^10^, which comprises 175,750,455 DNA nanoballs (≥1 UMIs) and 20,203 genes (≥100 UMIs tissue wide). While most analyses of such large data require binning or cell-segmentation preprocessing, Smoothie’s memory and runtime efficiency enabled direct smoothing over all DNA nanoballs in ∼1 hour on a standard CPU using ten parallel processes (**Fig. S3**). After Gaussian smoothing (20 μm Gaussian standard deviation), Smoothie identified 430,449 spatial gene correlations above a PCC threshold of 0.4, and 125,789 correlations above 0.5 (**Fig 3A, Data S3**). Clustering on the PCC > 0.4 network led to 272 gene modules, with 62 modules having 5+ genes (**Fig 3A, 3C**), far exceeding the 35 modules reported in the original publication. We visualized representative genes for the 16 largest spatial gene modules (**Fig 3B**) and generated a gene pattern atlas for all 89 gene modules with 3 or more genes (**Fig. S6, Data S4**). These modules correspond to specific organs, sub-organ regions, or cell types. For example, gene module 1 corresponds to nerve tissue (*Map2, Gap43, Gad1* markers), module 4 to muscle tissue (*Myod1, Myf5, Myog, Pax7* markers), module 5 to liver tissue (*Alb, Afp* markers), and module 6 to outer epidermal tissue (*Krt1, Krt10, Lor* markers).

**Figure 3.**
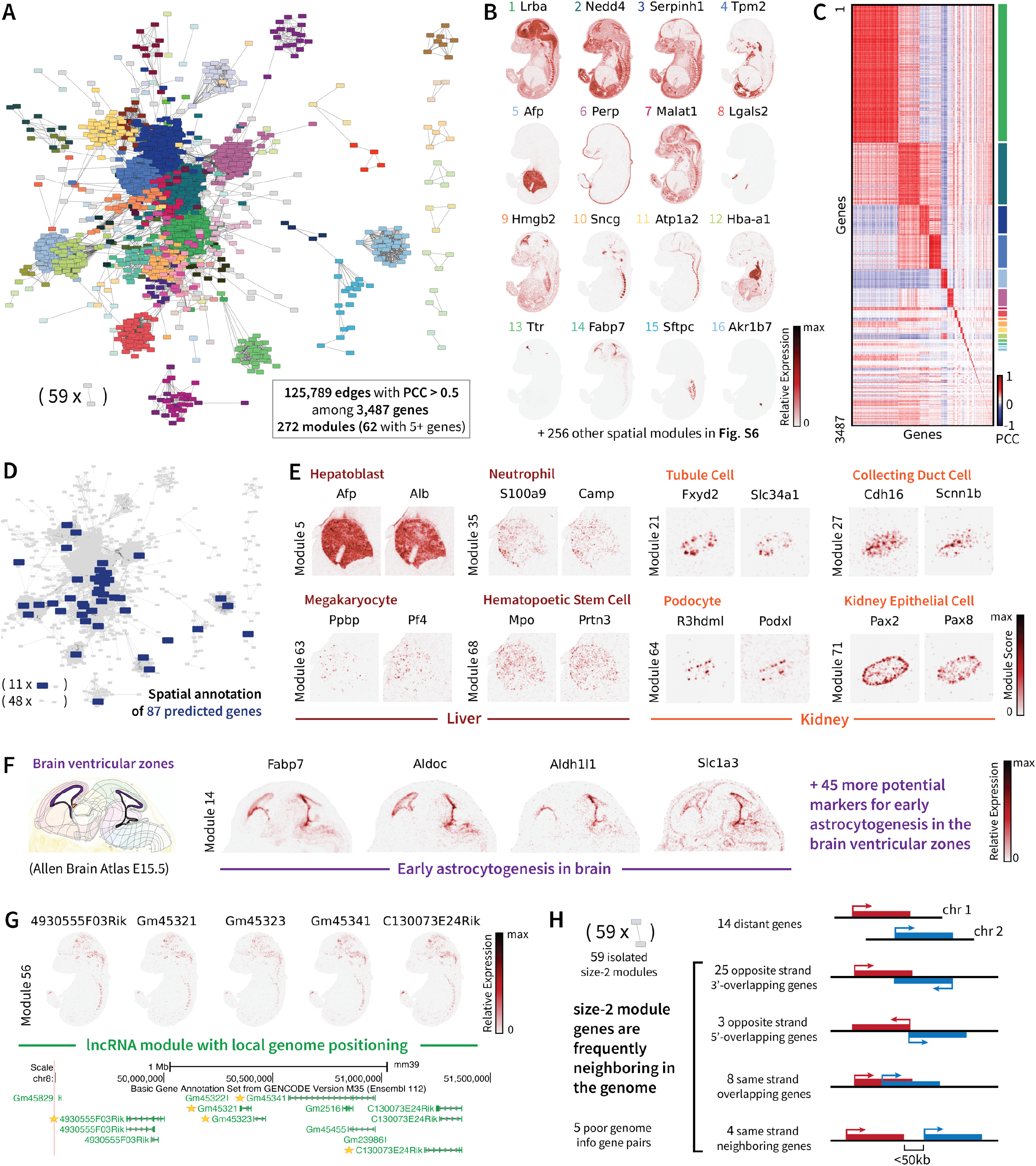
Smoothie reveals hundreds of spatial patterns in the developing E16.5 mouse embryo. **A**.The spatial gene correlation network produced from a Pearson correlation coefficient (PCC) cutoff of 0.5 finds 125,789 correlated gene pairs among 3,487 genes. Genes are colored by their assigned module. **B**. Example genes from the 16 largest Smoothie gene modules. Plots show relative gene expression after normalization and smoothing. **C**. The pairwise gene correlation matrix, with gene rows and columns ordered by assigned gene module. **D**. Smoothie spatially annotates 87 predicted or hypothetical mouse genes with no known gene function. **E**. Examples of sub-organ level classification of spatial gene modules in the liver (left) and the kidney (right). **F**. Smoothie identifies several known astrocyte marker genes and new potential astrocyte marker genes in the brain ventricular zone in early astrocytogenesis. **G**. Smoothie identifies a module of 5 lncRNAs that all reside within a 2Mb window of chromosome 8. **H**. Most size-2 Smoothie gene modules exhibit near or overlapping genome positioning.

Smoothie further delineated sub-organ cell type patterns in the liver, revealing marker genes for hepatoblasts, neutrophils, megakaryocytes, and hematopoietic stem cells in modules 5, 35, 63, and 68, respectively (**Fig 3E**). Similarly, in the kidney, Smoothie identified marker genes for tubule cells, collecting duct cells, podocytes, and kidney epithelial cells in clusters 21, 27, 64, and 71, respectively (**Fig 3E**). Examining module 14 in detail, we observed several astrocyte and radial glial cell marker genes in the brain’s ventricle zones at the key early stage in brain astrocytogenesis, prior to astrocyte migration and subtype differentiation^11–13^(**Fig 3F**). Module 14 contained 49 genes total, with a majority of such genes being new potential astrocyte markers during early phase astrocytogenesis (**Data S4**). In addition to tissue-specific programs, Smoothie identified spatial modules corresponding to widespread expression, spanning many to all organs, including modules 2, 3, 7, and others. In summary, the Smoothie-generated gene pattern atlas provides a thorough cataloging of genes based on their spatial expression, agreeing with known cell types and marker genes, while simultaneously revealing novel gene programs, new marker genes, and other biologically meaningful insights.

As we identified hundreds of thousands of strong spatial correlations, we asked how many of these gene pairs represented novel associations. To explore this, we focused on the 22,281 hypothetical or predicted (unannotated) mouse genes in the *NCBI Gene* database lacking geneRIF functional annotation (as of Nov. 1st 2024). From this set, 1,378 unannotated genes had ≥100 UMIs in the embryo dataset. Using a PCC > 0.4 cutoff, we detected 5,914 pairwise correlations involving these unannotated genes, allowing us to assign modules and infer spatial annotations for 87 of them based on known tissue regions, cell types, or gene pathways (**Fig 3D**). For example, we linked *Gm9947, Gm15533, Gm34081, Gm14635*, and *Gm14280* to muscle tissue (module 4), *Gm45941, Gm13629, Gm13425*, and *Gm12298* to *Sncg* neurons (module 10), *Gm26973* and *Gm19990* to lung tissue (module 15); *Gm7247* and *Gm14226* to the adrenal gland (module 16), and *Gm38505* and *Gm13889* to the subpallium brain region (module 29). We assigned modules to another 231 unannotated genes whose maximum PCC value fell between 0.2 and 0.4 by labeling each gene according to its highest-correlated neighbor in the PCC > 0.4 network. Overall, unannotated genes displayed diverse spatial patterns, appearing in 18 of the 20 largest modules and in 45 of 272 modules in total. If we also include genes with a maximum PCC of 0.2–0.4, the coverage extends to 72 modules. We provide a complete list of these new unannotated gene correlations in **Data S5** and the corresponding module assignments in **Data S6**.

Finally, we investigated the relationship between spatially correlated genes and genomic position, as genome organization and gene co-expression can be interconnected^14–17^. Using *UCSC Genome Browser*, first, we observed that spatial module 56 comprises 5 non-coding genes—4930555F03Rik, Gm45321, Gm45323, Gm45341, C130073E24Rik— that all fall within 2Mbp region on chromosome 8 (GENCODE *VM35*, **Fig 3G**)^18,19^. Genes 4930555F03Rik and Gm45321 fall on the positive strand, while the other three are negative strand genes. Moreover, there is a >300 kb gap between 4930555F03Rik and Gm45321, and a >200 kb gap between Gm45341 and C130073E24Rik, suggesting that co-expression in this cluster may be driven by local regulatory mechanisms. In contrast, genes in other spatial modules are not co-located; for instance, module 64, which contains multiple kidney podocyte markers, includes four genes each situated on a different chromosome. We also examined the genomic locations of 59 gene pairs that formed “isolated” size-2 spatial modules in the network (i.e., each pair had a PCC > 0.5, and neither gene in a pair had a PCC > 0.4 with any other gene). Of these 59 pairs, 14 had genes situated on separate chromosomes or ≥15 Mb apart on the same chromosome, 44 comprised “overlapping” or closely spaced genes (<50 kb apart) on the same chromosome, and 1 pair was discarded because one member (Gm26776) lacked a known genomic locus. Closer inspection of the 44 pairs within 50 kb (using GENCODE *VM31* and *VM35* annotations) showed that 25 span opposite strands with 5' overlap or proximity, 3 share opposite strands with 3' overlap or proximity, 8 overlap on the same strand (potentially multi-mappers), 4 are consecutive neighbors on the same strand, and 4 pairs involved a chromosome accession number rather than a resolvable gene ID (**Fig. 3H**). This shows how genes located near each other in the genome are often enriched in these size-2 modules, with many pairs overlapping on opposite strands.

### Smoothie characterizes the spatial patterns of the Hox family and Slc transporter family in the E16.5 embryo

We next used Smoothie to examine the spatial organization of gene sets of interest in the E16.5 mouse embryo, focusing first on the well-characterized Hox gene family. Hox genes are expressed during the development of almost all animals and exhibit defined anterior–posterior expression domains^20^. Out of the 39 mouse Hox genes, 33 showed non-random spatial correlations (along with 3 antisense transcripts), while the remaining 6 were too sparsely expressed to characterize. The Hox gene correlation network revealed that anterior Hox gene types 2– 5 cluster on one side, whereas posterior Hox gene types 10–13 occupy the opposite side of the network (PCC > 0.2, **Fig. 4A**). Traversing the Hox network from left to right, we can visualize how the gene patterns change in their expression along the anterior-posterior axis, validating our spatial network approach (**Fig. 4B**).

**Figure 4.**
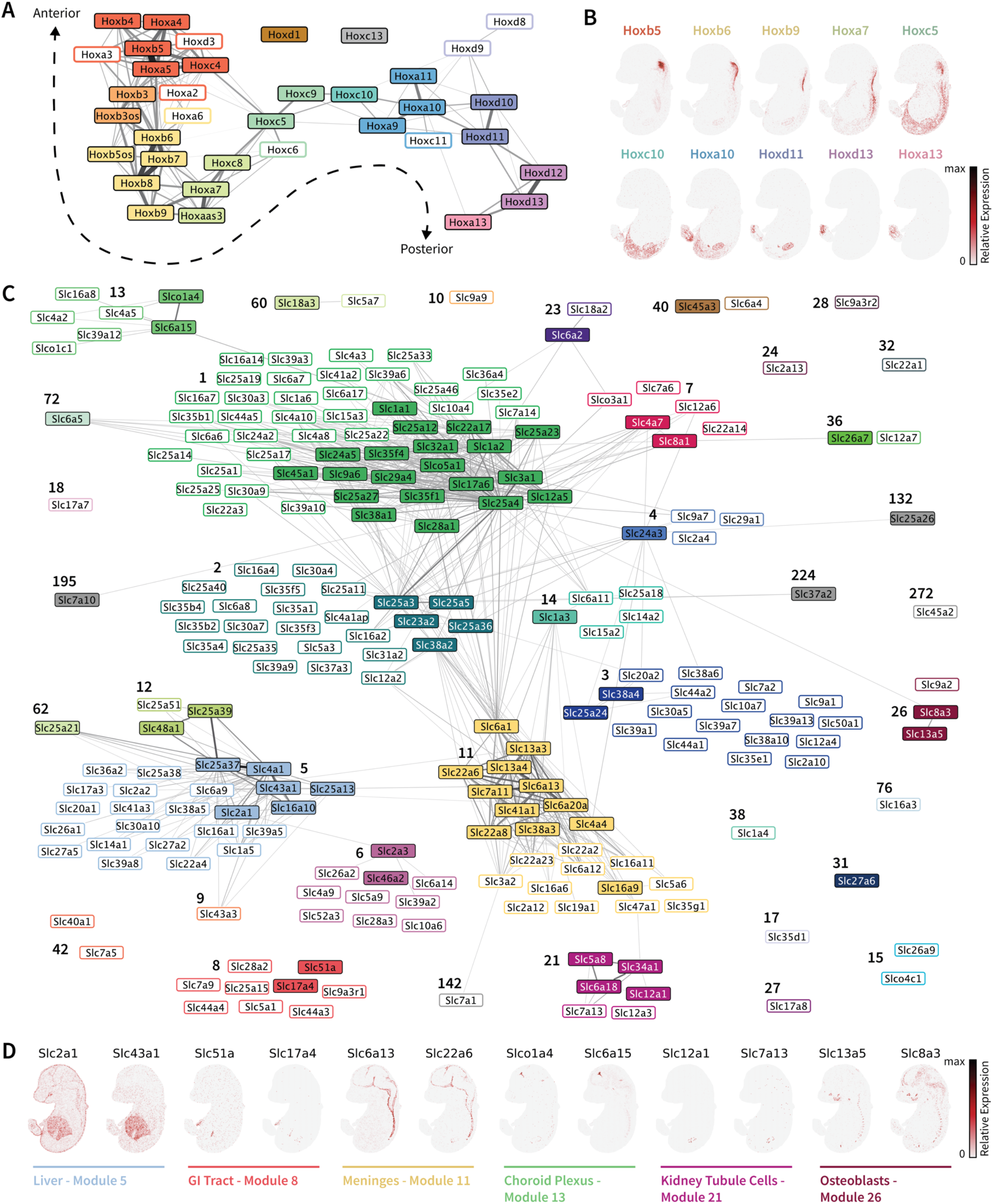
Smoothie comprehensively summarizes the spatial pattern distribution of the Hox gene family and the solute carrier (Slc) gene family in the E16.5 mouse embryo. **A**.The spatial Hox gene correlation network reveals the known anterior to posterior axis of expression. Edge width and darkness reflect the Pearson correlation coefficient (PCC); genes are colored by module; solid-colored genes have at least 1 correlation with PCC > 0.4; and hollow genes have no correlation with PCC > 0.4 but at least 1 correlation with PCC > 0.2. **B**. Example Hox genes moving across the network from the anterior to posterior side. Plots show relative gene expression after normalization and smoothing. **C**. The spatial Slc gene family correlation network reveals 38 modules of Slc genes. Edge and node specifications are identical to those in Fig. 4A. **D**. Example Slc genes across different spatial modules. Plots show relative gene expression after normalization and smoothing.

We then analyzed the spatial distribution of solute carrier (Slc) membrane transport proteins (**Fig. 4C**). The Slc superfamily consists of over 400 genes across 65 families, each responsible for transporting specific ions, hormones, or other solutes^21–24^. Over 100 Slc genes have been implicated in human diseases, and their roles in molecular transport make them attractive targets for tissue-specific drug delivery^21–23^. Of the 356 Slc genes with 100+ UMIs in the embryo dataset, the PCC > 0.4 network organized 73 Slc genes into modules. Using a more permissive cutoff (PCC > 0.2), an additional 152 Slc genes were assigned to modules in this network. In total, 225 Slc genes were thus organized into 38 modules (**Fig. 4C**). These modules highlight Slc gene groups predominantly expressed in specific embryonic regions, such as the CNS (52 genes in module 1), liver (24 genes in module 5), meninges (23 genes in module 11), epidermis (10 genes in module 6), GI tract (9 genes in module 8), choroid plexus (7 genes in module 13), kidney tubule cells (6 genes in module 21), osteoblasts (3 genes in module 26), and others (**Fig. 4D; Data S7**). Notably, many of these tissues are associated with fluid filtration— liver (blood), meninges (blood-brain barrier), choroid plexus (cerebrospinal fluid), GI tract (bile), and kidney tubule cells (blood). Other Slc genes show broad expression across multiple organs (e.g. modules 2, 3, and 7), potentially indicating housekeeping functions or expression in widespread cell types. This analysis aligns with a recent finding that no single Slc family is restricted to one tissue region^21^. Nevertheless, we detect enrichment of Slc6, Slc13, Slc16, and Slc22 family genes in the meninges (module 11), whereas certain Slc families (e.g., Slc25, Slc38) exhibit more pervasive expression patterns.

### Smoothie presents a new way to analyze gene modules across multiple spatial experiments

Last, we show how Smoothie can integrate spatial transcriptomes from multiple samples to ***i)*** find gene modules spanning multiple conditions, and ***ii)*** quantify gene pattern stability or variability across conditions. To measure multi-dataset gene modules, we concatenated the normalized, smoothed gene surfaces of individual datasets into a combined dataset prior to pairwise gene correlation calculation (**Methods**). Applying this approach to the MOSTA mouse embryogenesis data at E13.5, E14.5, E15.5, and E16.5 revealed 782,486 correlated gene pairs (PCC > 0.4) among 6,419 genes, organized in 298 gene modules (69 modules with ≥ 5 genes; **Fig. 5A; Data S8, Data S9**)^10^. We used Enrichr to annotate modules by cell-type enrichment in several popular scRNA-seq atlases (**Fig. 5B**)^25–32^. Some examples include module 4 (skeletal muscle, 274 genes), module 14 (chondrocytes, 44 genes), module 25 (lung and intestine smooth muscle, 25 genes, module 37 (Type II pneumocyte, 12 genes), module 65 (all *Zic* family genes), and module 92 (mesothelial cell layer, 3 genes), with all 111 modules (size ≥ 3) visualized in **Fig. S7**.

**Figure 5.**
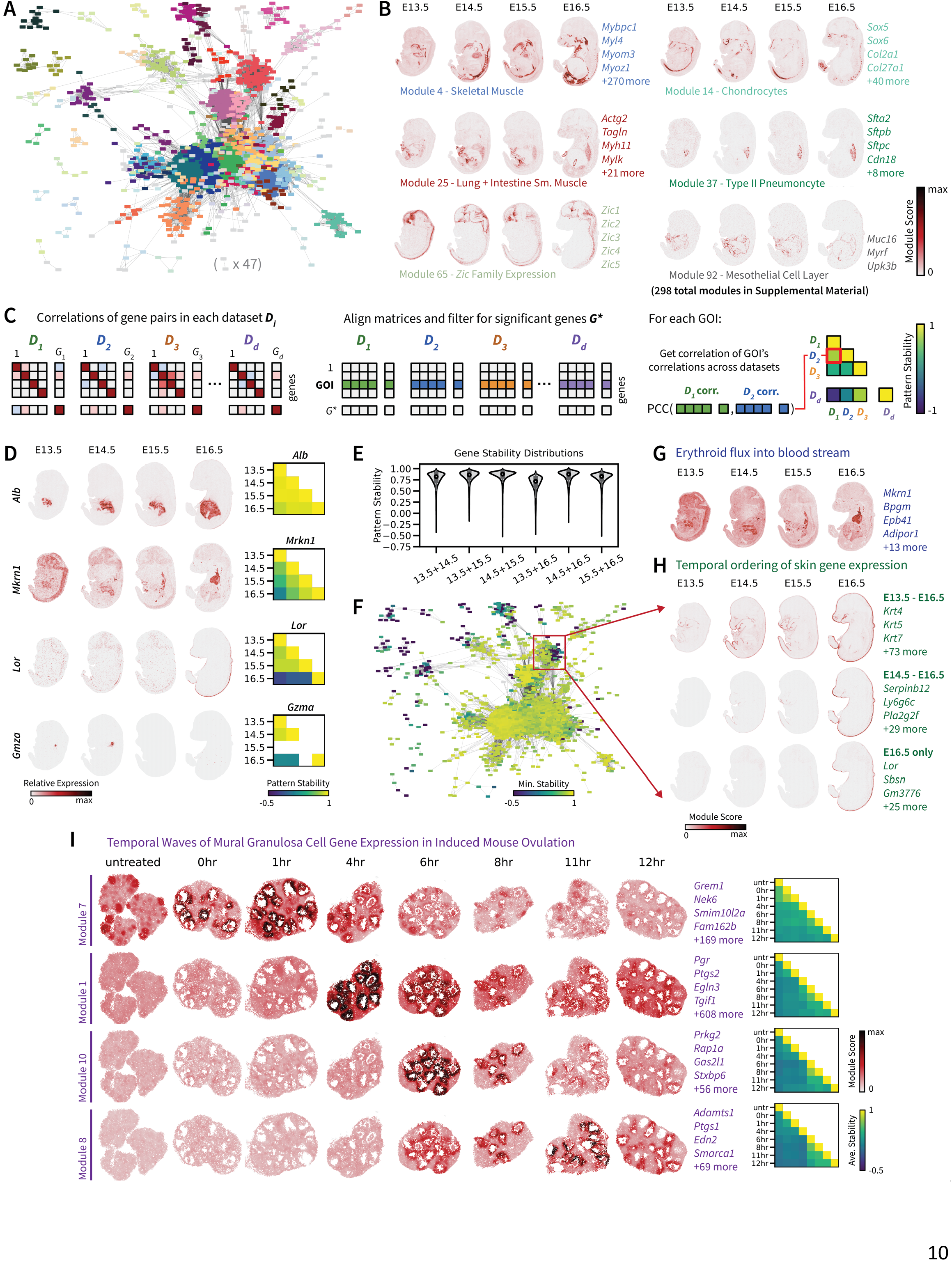
Smoothie finds multi dataset gene modules and ranks gene patterns from stable to dynamic. **A**.The gene correlation network created by comparing the joint gene patterns of all E13.5-E16.5 mouse embryos. **B**. Examples of gene patterns and marker genes for modules in the network from 5A. There are 298 modules with 2+ genes, and 69 modules with 5+ genes. Plots show the average 0 to 1 scaled genes for the module. **C**. Overview of the second-order correlation logic used to rank gene patterns at stable or dynamic across spatial datasets of mismatched (*x, y*) coordinates. In short, intra-dataset pairwise gene correlations are used as gene embeddings that can be compared across datasets. **D**. Examples of a stable gene pattern—liver marker *Alb*—and three dynamic gene patterns—*Mkrn1, Lor*, and *Gmza*. **E**. The distributions of gene stabilities values across each dataset pair. Genes that are missing in one or the other dataset are not included in these distributions. **F**. The spatial correlation network from 5A with gene stability node coloring reveals modules of dynamic gene patterns. Missing genes take the value of -1 for their stability in this network. **G**. Example of a dynamic gene module containing 17 genes with several erythroid cell marker genes. **H**. Gene stability analyses find temporal skin expression patterns. Some genes exhibit stable skin expression from E13.5-E16.5; some genes show skin expression from E14.5-E16.5; and some genes are only expressed at E16.5. **I**. Smoothie identifies temporally restricted waves of gene expression in the antral mural granulosa cells throughout 0-12 hour induced ovulation. For panel I only, the module score 0-max scale is shared across datasets.

To explore variation in gene expression across these four embryonic stages, we first generated pairwise gene correlation matrices for each dataset independently, capturing 7,438 genes with at least one significant correlation (PCC > 0.4) in any dataset. We then quantified second-order correlations to ask: “Does gene *A* maintain similar correlation profiles among other genes in dataset *D*_1_ and dataset *D*_2_ ?” (**Methods**). This second order correlation quantifies gene *A*’s pattern stability across datasets (**Fig 5C**). As shown in **Figure 5E**, most genes had high stability (median values 0.7–0.9) across dataset pairs. For instance, the *Alb* liver marker remained consistently expressed, whereas *Mkrn1, Lor*, and *Gmza* varied significantly—*Mkrn1* expanded into arteries and heart by E16.5, *Lor* activated only at E16.5 in epidermal tissue, and *Gmza* disappeared at later stages because those tissue slices lacked the thymus (**Fig. 5D**). Combining the multi-dataset correlation network and the gene stability scores allowed us to identify temporally variable gene modules (**Fig. 5F**). For example, module 19 included numerous erythroid markers (e.g. *Mkrn1, Bpgm, Epb41, Adipor1*) that shifted from liver to blood vessel expression between E13.5 and E16.5, perhaps reflecting changes in erythroid cell abundance and location during embryogenesis^33^. Module 8 (epidermis) revealed temporal waves of gene activation—76 genes remained stable from E13.5–E16.5, 32 became activated from E14.5 onward, and 28 appeared only by E16.5 (including *Lor, Sbsn, Lipk*; **Fig. 5G, Data S10**), revealing new insights on the periderm’s maturation^34,35^.

Finally, we applied Smoothie to Slide-seq V2 spatial transcriptomics data of the mouse ovary collected at 8 distinct time point in the scope of hormone-induced ovulation (0-12h post induction)^36,37^. Because each sample’s follicle arrangement and stage in the ovary differ, integrating the data via coordinate warping or optimal transport alignment is both impractical and error-prone in this case. Instead, Smoothie identified 131,284 correlations (PCC > 0.45) among 2,689 genes, yielding 70 modules (33 with ≥5 genes) (**Data S11, Data S12; Fig. S8**). Several Smoothie modules agreed with the spatial cell type labels in the original paper^37^. For instance, module 2 (523 genes) corresponded to the cumulus oocyte complex (COC), capturing known markers (*Areg, Btc, Bmp15, Dazl, Ereg, Gdf9*) as well as new candidate regulators, including 15 Fbxw and 3 Fbxo genes implicated in SCF ubiquitin ligase function^38^. It has been shown that *Fbxw7* and *Fbxw24* knockout mice exhibit severe fertility defects^39,40^; however, the overall connection of the Fbxw and Fbxo gene families to the COC during oogenesis is understudied.

Further, we observed large temporal changes in gene expression within the antral mural granulosa cells (MGC) annotated in the original paper^37^. For example, Module 7 contained 173 genes with shared expression at 0–1 hours, Module 1 had 612 genes with shared expression at 4 hours, Module 10 included 60 genes expressed at 6–8 hours, and Module 8 had 73 genes expressed between 6–12 hours (**Fig. 5I**). These modules reveal wave-like changes in antral MGC gene programs over the 12-hour ovulation process, involving numerous regulatory factors and reported ovulation genes—such as *Pgr* and *Ptgs2* at 4 hours, and *Adamts1* and *Edn2* between 6–12 hours^41–44^. Plotting the average gene pattern stabilities for each of these modules supports the antral MGC expression shifts too (**Fig. 5I**). Collectively, these examples illustrate Smoothie’s utility for identifying, quantifying, and visualizing gene expression variation across diverse tissues.

## DISCUSSION

In this work, we introduced Smoothie, a computationally efficient method for measuring spatial gene correlation networks from denoised spatial transcriptomics data. Smoothie’s use of Gaussian smoothing addresses data sparsity and noise, which expands the pool of analyzable genes in downstream correlation analyses. The ability to measure spatial correlations and assign gene modules for a larger number of genes (up to 5,000 to 10,000 genes for some datasets) increases the potential for finding new biology. This sets Smoothie apart as an effective spatial gene module detection and biological discovery tool, even for challenging, low-UMI datasets.

Smoothie’s potential for biological discovery comes from the abstraction of raw spatial transcriptomics data into pairwise gene correlations and spatial gene modules. The modules capture many facets of tissue biology. For instance, we found that many spatial modules map onto single or closely related cell types (e.g., granule vs. Purkinje neurons in the cerebellum, or podocyte vs. tubule cells in the kidney), which can reveal new potential marker genes in those modules. Another major use case for Smoothie is gene characterization, which could be particularly impactful for studies of non-model organisms, where gene and functional annotations are often sparse. Even in the well-annotated mouse model and using data from just one experiment, we were able to spatially annotate hundreds of unknown genes, which gives confidence that this approach will be effective in understudied non-model organisms. Interestingly, we also observed modules whose genes cluster by genome position. Integrating Smoothie’s networks with motif analysis or Hi-C chromatin interaction data could further illuminate how local genome organization influences spatial transcript patterns and gene regulation. Finally,

Smoothie also enables targeted investigations of particular genes, families, or pathways. For example, our analyses of the Hox and Slc gene families highlight the possibility to zoom in on a gene subset of interest, identifying submodules within correlation network. Gene set analyses can combine a targeted subset network (for clarity) with the broader correlation network (for context), both of which Smoothie supports natively.

A further major application area for Smoothie is found in multi-sample analysis and data integration. While numerous solutions have been proposed for integration of single-cell transcriptomic data, spatial transcriptomics has few robust solutions to integrate data across replicates, time points, conditions, or species^45,46^. Morphing of multiple tissues to match spatial patterns has been proposed, but this is error prone. Single-cell segmentation followed by integration is another possibility, but it is sensitive to any imperfections in segmentation, which can be substantial. Here, we suggest that the abstraction of the raw data enabled by measuring gene correlation networks provides a compelling avenue for data integration. We propose and implement two methods: ***i)*** measuring gene correlations of all smoothed datasets together to find multi-dataset gene modules, and ***ii)*** measuring second-order gene correlations across datasets to compare gene patterns across datasets. Such approaches to data integration can enable efficient comparison of spatially resolved transcriptomes from multiple tissues, and time points. In addition, cross-species integration of spatial gene network data is an avenue for future investigation that can be impactful. Because measuring gene networks does not require consideration of the tissue architecture or biology, integration of spatial gene correlation networks may form the basis for the construction of large-scale spatial transcriptomic atlases of organ systems and organisms, akin to what has been accomplished with single-cell transcriptomic atlases.

We have shown that Smoothie is both effective and scalable on platforms ranging from Slide-seq (10 μm resolution) to Stereo-seq (0.5 μm resolution), providing confidence in its utility for other ST technologies—such as OpenST (0.6 μm), Visium-HD (2 μm), and beyond^6,10,36,47^. Further, we foresee that Smoothie’s framework is adaptable to a range of other applications. Briefly, Smoothie could find correlations between spatial distributions of any data modality, e.g. protein expression, chromatin accessibility, metabolite levels, etc., and can thereby support multi-omics data integration. Furthermore, as the Gaussian smoothing kernel can be adjusted to account for three-dimensional distances, 3D spatial transcriptomic datasets could be analyzed via Smoothie. Last, Smoothie’s ability to recover biological signals in noisy datasets will make it useful for applications where tissue quality can be poor, including clinical applications. As spatial omics technologies improve in resolution and scope, we anticipate Smoothie’s utility will only grow, enabling adoption of a “forward spatial transcriptomics paradigm”—one in which data-driven, network-based approaches reveal novel biology rapidly across samples of different phenotypes or conditions.

## METHODS

### Single Dataset Smoothie Pipeline

#### Gaussian smoothing

Smoothie first performs Gaussian smoothing on each gene in the normalized spatial count matrix *X* to reduce data noise and sparsity. This process entails moving a 2D Gaussian distribution around the spatial data and taking a weighted average of the gene values under the distribution to generate the smoothed gene surface. The standard deviation *σ* of the 2D Gaussian specifies the degree of smoothing (**Fig. S1**). The smoothing radius *ρ* is the range of points extending from the center of the Gaussian to include in the smoothing process. To avoid extensive and unnecessary compute power, the radius *ρ* is set to be 3*σ* in the analysis, which retains 99.7% of the volume under the Gaussian. There is an additional variable *τ* that specifies the minimum number of coordinates required within the Gaussian’s radius for smoothing to occur. This prevents dividing by a small sum of values which can lead to noisy or inflated smoothed gene values in some cases. The effect of variable *τ* is similar to the computer vision and deep learning concept of applying convolution layers without padding, slightly shrinking the dataset in edge regions of the tissue to ensure meaningful results. Altogether, a Gaussian centered at a given spatial coordinate (*x, y*) generates the smoothed gene value 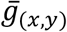 as follows:

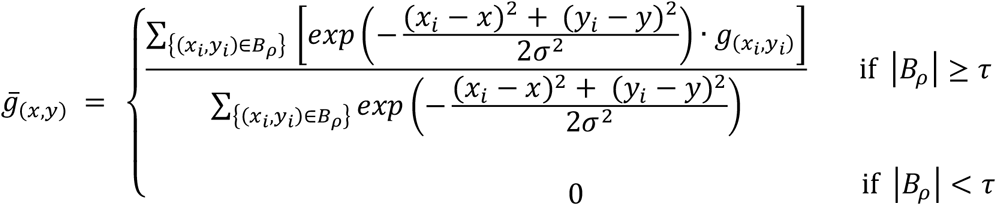

where, 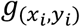 is the unsmoothed gene value at a given point (*x*_*i*_, *y*_*i*_) and *B*_*ρ*_ is the set of points that fall within the radius *ρ* from the point (*x, y*). The denominator contains the sum of the Gaussian values at each point within radius *ρ* to normalize for the non-uniform densities of spatial datapoints found in many spatial transcriptomic experiments. The accuracy of smoothing across genes is validated by plotting the total gene value sums for each gene before and after smoothing, revealing a near perfectly correlated line (**Fig. S1**).

For scalability to very large datasets, Smoothie has two implementations of Gaussian smoothing – in-place and grid-based smoothing. In-place smoothing centers a Gaussian at every single spatial location in the data and generates a smoothed value for each spatial location. Grid-based smoothing instead imposes a hexagonal grid onto the original data with stride *s* distance between neighboring grid points and then smooths only at the imposed grid points. For binned or cell-segmented data, in-place smoothing is effective and standard. Grid-based smoothing is designed for smoothing directly on the higher-resolution platforms’ data output, such as the 100s of millions of Stereo-seq DNA nanoballs. In the process of smoothing, the sparse gene count matrix is converted into a dense matrix, so grid-based smoothing is required for matrices with several millions of spatial locations.

To summarize, the input parameters to Smoothie consist of the spatial gene count matrix *X*, Gaussian standard deviation *σ*, Gaussian radius *ρ*, Gaussian coordinate threshold τ, a Boolean value *b* to specify in-place or grid-based smoothing, and the hexagonal grid stride *s*for grid-based smoothing only. In practice, these parameter choices can be reduced to be in terms of just *X* and *σ* for simplicity and effective results:

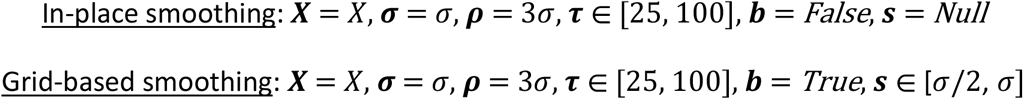

Effective choices for *σ* range from 10-50 µm and vary slightly across different spatial platforms. For Slide-seq, 30 µm is the default *σ* value, and for raw DNA nanoball Stereo-seq data, 20 µm is the default *σ* value. To validate the range of effective choices of *σ*, correlations of known spatially co-expressed cell type markers can be compared with correlations of random gene pairs across different levels of smoothing (**Fig. S1**). The choice of τdepends on data sparsity following data preprocessing as well as the choice of *σ* . A *τ* value of 25-100 is suitable for most datasets.

#### Gaussian smoothing scalable implementation

Since all genes in a spatial dataset share the same spatial coordinates, Smoothie can generate the smoothed values for all genes in parallel with matrix multiplication. Further, since generating smoothed gene values only requires information on the points within radius ρfrom the center of the Gaussian, parallelization allows multiple processes to smooth different regions of the dataset simultaneously. The runtime complexity for smoothing is *O*(*N*′.*G.ρ*^2^ .log(*N*)/*P*), where *N*is the number of spatial spots in the input, *N*′ is the number of spatial spots in the smoothed output, *G* is the number of genes, and *P* is the number of parallel processes. Smoothie benefits from efficient KD-tree queries that introduce only a logarithmic dependence on *N*. For in-place smoothing, *N*′= *N*, and for grid-based smoothing *N*′<N. Memory complexity of the smoothed output is *O*(*N*′.*G*), and a chunk count parameter *C* allows for efficient smoothing of smaller data segments, minimizing memory overhead during multi-processing. This combination of parallelization and chunking enables Smoothie to achieve fast, scalable runtimes while maintaining low memory requirements, even for large Stereo-seq datasets with hundreds of millions of DNA nanoballs.

#### Creating the gene correlation matrix

Smoothie constructs the pairwise gene correlation matrix *R* for all gene pairs in parallel with its own efficient matrix multiplication function. For a smoothed count matrix with *N* ′spatial spots and *G* genes, efficient computation of the (*G* choose 2) gene pair correlation values is necessary, as *G* typically ranges between 10,000 to 30,000 for unbiased sequencing-based spatial platforms, and *N*′ can be very large too, ranging from 10,000 to 1,000,000+ smoothed cells or spatial spots. The runtime complexity for constructing the pairwise gene correlation matrix *R* is *O* (*N* ′.*G* ^2^), with memory complexity *O* (*G* ^2^).

#### Permutation tests to find significant gene correlations

With the gene correlation matrix *R*, Smoothie determines which spatial gene correlations are significant to include in downstream analyses. While each gene pair’s Pearson correlation coefficient (PCC) is accompanied by a *p* value, the smoothing process artificially introduces a positive offset in correlation, preventing the *p* values from accurately defining a significance cutoff, even after FDR-correction. Instead, Smoothie uses a permutation test to randomly shuffle the spatial coordinates of the input count matrix *X*, generating *X*_*shuffled*_ . Smoothie performs Gaussian smoothing with the same input parameters on both *X* and *X*_*shuffled*_ and generates pairwise gene correlation matrices *R* and *R*_*shuffled*_ . From the distribution of all randomized gene pair correlations in *R*_*shuffled*_, the 99.9^th^ percentile cutoff *PCC*_99.9_ is used to select for significant, non-random gene correlations in the smoothed true dataset gene correlation distribution (**Fig. S1**). There is also an option to shuffle cell-sized bins of spatial coordinates for subcellular resolution spatial platforms like Stereo-seq. Non-binned shuffling of subcellular coordinates would be a poor null hypothesis as there is co-expression of genes occurring within any given cell.

#### Creating and clustering the gene correlation network

Let *X* be a spatial count matrix with gene set 𝒢 = {*G*_1_, *G*_2_,... *G*_*G*_}; let *R* be the pairwise gene correlation matrix derived from a smoothed version of *X* ; and let *PCC* * be a hard threshold for the matrix *R* . The weighted, undirected gene correlation network *W* with node set *V* and edge set *E* is defined as:

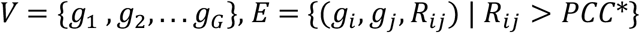

The network *W* is clustered into gene modules using the Infomap random-walk clustering algorithm implemented in the Python package *igraph* ^7,48^. Before clustering, Smoothie applies an additional soft thresholding transformation *f*_*β*_ on the edge weights after applying the hard threshold *PCC* *. The transformation *f*_*β*_ linearly rescales edge weights from the interval [*PCC* *,1] to [0,1] and then raises the values to a power *β* ≥ 1. The transformation *f*_*β*_ is monotonic and serves to emphasize stronger correlations while suppressing weaker ones. Higher values of *β* generally lead to a more modular network by making strong correlations more dominant in clustering, whereas lower values of *β* produce fewer, larger clusters. The choice of *PCC* * will change the size of the output network. Higher *PCC* * values generate smaller networks with higher average correlation values, whereas smaller *PCC* * values generate larger networks with lower average correlation values. A discussion of thresholding strategies and scale-free topography of co-expression networks can be found in this paper^1^. Smoothie includes a function to measure the scale-free topography index of the network across different choices of *PCC* * and *β* for consideration in network construction.

#### Visualizing the correlation network

Smoothie’s network output files can be imported into Cytoscape for visualization^49^. The “Prefuse Force Directed Layout” setting positions nodes in the network, and nodes may then be colored by their gene module assignment, chromosome number, binary values specifying gene set membership, or other variables.

### Multiple Dataset Smoothie Pipeline

#### Gaussian smoothing

Given a set of *d* spatial count matrices to integrate, the first step after normalization is to smooth each spatial count matrix in the set with equal smoothing parameters, as described above. Let {*X*_1_, *X*_2_,..., *X*_*d*_} be the set of smoothed spatial count matrices, where each *X*_*i*_ has *N*_*i*_ spatial spots and *G*_*i*_ genes after smoothing.

#### Creating and clustering the gene correlation network for multiple datasets

The set of smoothed count matrices can be concatenated across spatial coordinates to create one smoothed count matrix *X*_*concat*_ with *N*_*concat*_ spatial spots (rows) and *G*_*concat*_ genes (columns). Here *N*_*concat*_ is the total number of points across all matrices and *G*_*concat*_ is the number of unique genes across all matrices. The creation of *X*_*concat*_ is such that gene columns are aligned across datasets during concatenation, and if a gene is missing in a given dataset, that dataset’s portion of the gene column is filled with zeros. With *X*_*concat*_, Smoothie can then create the pairwise gene correlation matrix *R* as before using a spatially permuted version of *X*_*concat*_ to determine the *PCC*_99.9_ threshold of *R*_*shuffled*_ . After creating the weighted network *W* using matrix *R* and threshold *PCC* * ≥ *PCC*_99.9_ as explained before, Infomap clustering reveals spatial gene modules and the network can be visualized for interpretation.

#### Cross dataset gene pattern embedding comparisons

Here, we propose a new approach to measure the similarity of two genes’ spatial expression patterns across different spatial datasets. Given a set of smoothed spatial gene count matrices {*X*_1_, *X*_2_,..., *X*_*d*_}, where each *X*_*i*_ has *N*_*i*_ spatial spots and *G*_*i*_ genes, these datasets have different tissue morphologies and different (*x, y*) coordinates. Thus, the smoothed gene surfaces between two distinct count matrices cannot be directly compared via Pearson correlation. Instead, Smoothie leverages a second-order correlation approach. After generating a gene pair correlation matrix *R*_*i*_ for each smoothed count matrix 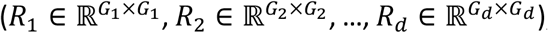, Smoothie creates aligned versions of the correlation matrices for each dataset 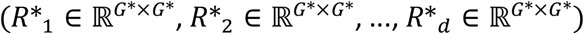 where 1) row *j* and column *j* are the same gene across all *R* ^*^_*i*_ and 2) *G* ^*^ is the magnitude of the gene set 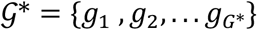 of genes that have a Pearson correlation coefficient above a threshold *PCC* ^*^ in at least 1 dataset. Genes that are missing (< 100 UMIs) in one dataset but significant in another dataset have their missing gene’s correlations filled with Null values. The threshold *PCC* ^*^ is determined considering the *PCC*_99.9_ values determined with permutation tests for each individual dataset.

Genes in 𝒢^*^ can then be compared across datasets by measuring the correlation of any two rows from the aligned matrices *R* ^*^_1_, *R* ^*^_2_, …, *R* ^*^_*d*_ . In other words, for each dataset, a given gene *G* in 𝒢^*^ has a gene embedding row vector 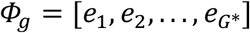 where *e*_*i*_ is the correlation value of gene *G* and gene *i*, and these embedding vectors are comparable across datasets. Prior to comparing embedding vectors, the Fisher’s Z transformation is applied to all correlation values across all *R* *_*i*_ to stabilize the variance of the correlation distributions. If two gene embeddings are similar across two datasets, then those genes have similar spatial expression patterns, whereas dissimilar embeddings indicate variable spatial expression patterns. The formula for measuring gene similarity between gene *A* in *R* *_1_ and gene *B* in *R* *_2_ is simply the Pearson correlation coefficient of the two gene embedding vectors 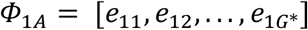 and 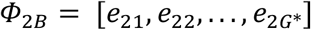, using only embedding features with finite values in both vectors in the calculation:

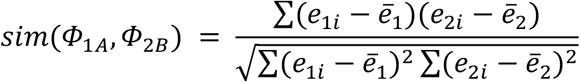

This logic relies on two key assumptions:

##### 1. Gene embeddings are comparable across datasets

Comparing genes across datasets using their intra-dataset gene correlations requires that a majority of genes have comparable or “stable” expression patterns across datasets (i.e. most gene embedding features are comparable). Each comparable embedding feature contributes to the success of the second order correlation approach, whereas unaligned embedding features add noise to the calculation. By using only the 𝒢^*^ significantly correlated genes for this analysis, Smoothie filters out genes with less structured and more noisy patterns, which increases the ratio of stable to unstable genes. In some cases, the corresponding embedding features for known unstable genes may be filtered out prior to analysis.

##### 2. Gene embeddings adequately cover the complexity of gene patterns

Gene embeddings must also span the complexity of gene patterns present in the tissue slices with near equal pattern weights. Here, embeddings are designed as gene correlations with all other genes in 𝒢^*^, maximizing pattern diversity without having to add any synthetic features. Since some patterns contain more genes than others, a parameter *e*_*max*_ limits the number of genes per module used as embedding features. If a module exceeds *e*_*max*_, its features are randomly down sampled. After down sampling, each matrix *R* ^*^_*i*_ retains the same number of genes to compare but has fewer embedding features for cross-dataset comparisons.

Software Requirements

Smoothie was implemented using Python 3.10.12, along with the following packages: anndata (v0.9.2)^50^, igraph (v0.10.6)^48^, numpy (v1.23.5)^51^, pandas (v1.5.3)^52^, scanpy (v1.9.3)^53^, and scipy (v1.9.3)^54^.

Data resources, preprocessing, and normalization

#### Mouse cerebellum data, Slide-seq

The Slide-seq Puck_180819_12 dataset .h5ad file was obtained from the Hotspot github page https://hotspot.readthedocs.io/en/latest/Spatial_Tutorial.html. This data is also available from the original paper here https://singlecell.broadinstitute.org/single_cell/study/SCP354/slide-seq-study#study-summary^6^. Genes expressed at least 50 different spatial locations were kept for the analysis. Data was normalized with counts per thousand (CPT) normalization and then log1p normalization.

#### Mouse embryo development data, Stereo-seq

The raw DNA nanoball datasets E13.5_E1S1_GEM_bin1.tsv.gz, E14.5_E1S1_GEM_bin1.tsv.gz, E15.5_E1S1_GEM_bin1.tsv.gz, and E16.5_E1S1_GEM_bin1.tsv.gz were obtained from https://db.cngb.org/stomics/mosta/download/ and converted to .h5ad files^10^. DNA nanoballs with 1 or more UMIs and genes with at least 100 total UMIs were kept. Data was normalized with only log1p normalization, as CPT normalization on DNA nanoballs with 1 to a few UMIs is nonsensical.

#### Mouse ovary timecourse data, Slide-seq V2

Sequencing data is available with GEO number GSE240271 and spatial barcode files are listed in https://github.com/madhavmantri/mouse_ovulation^37^. Spatial spots with at least 100 UMIs and genes with at least 50 total UMIs were kept. Data was normalized with CPT normalization and then log1p normalization.

### Smoothie Implementation

#### Slide-seq mouse cerebellum

In-place gaussian smoothing was performed on Slide-seq beads with gaussian standard deviation ***σ*** = 30 μm, gaussian radius ***ρ*** = 90 μm, and minimum coordinates under gaussian ***τ*** = 100. The PCC distribution of permuted spatial coordinate gene correlations found a *PCC*_99.9_ cutoff of 0.2165. The correlation network was built with a ***PCC***^*^ >0.35, and the clustering power ***β*** = 4 was used for gene module identification.

#### Stereo-seq E16.5 mouse embryo

Grid-based gaussian smoothing was performed on Stereo-seq DNA nanoballs with gaussian standard deviation ***σ*** = 20 μm, gaussian radius ***ρ*** = 60 μm, minimum coordinates under gaussian ***τ*** = 100, and hexagonal grid stride ***s*** = 20 μm. The PCC distribution of permuted spatial coordinate gene correlations found a *PCC*_99.9_ cutoff of 0.1098. The correlation network was built with a ***PCC***\* >0.4, and the clustering power ***β*** = 2 was used for gene module identification.

#### Stereo-seq E13.5 to E16.5 mouse embryo time course

Grid-based gaussian smoothing was performed on DNA nanoballs of each embryo dataset—E13.5, E14.5, E15.5, E16.5—with identical smoothing parameters of gaussian standard deviation ***σ*** = 20 μm, gaussian radius ***ρ*** = 60 μm, minimum coordinates under gaussian ***τ*** = 100, and stride ***s*** = 20 μm. For multi-dataset module detection, the *PCC*_99.9_ cutoff generated from the combined permuted datasets was 0.406. The combined dataset correlation network was built with a ***PCC***^*^ >0.4, and the clustering power ***β*** = 4 was used for gene module identification. For gene embedding comparison analyses, the maximum *PCC*_99.9_ cutoff generated among the permuted individual datasets was 0.116. The set of 𝒢* genes for embedding comparison analysis were selected with cutoff ***PCC***^*^ >0.4, and the maximum feature embeddings per module was capped at ***e***_***max***_ = 25.

#### Slide-seq V2 mouse ovaries time course

In-place gaussian smoothing was performed on the Slide-seq V2 beads of each ovary dataset—untreated, 0hr, 1hr, 4hr, 6hr, 8hr, 11hr, and 12hr—with identical smoothing parameters of gaussian standard deviation ***σ*** = 30 μm, gaussian radius ***ρ*** = 90 μm, and minimum gaussian coordinates ***τ*** = 25. For multi-dataset module detection, the *PCC*_99.9_ cutoff generated from the combined permuted datasets was 0.420507. The combined dataset correlation network was built with a ***PCC***^*^ >0.45, and the clustering power ***β*** = 4 was used for gene module identification. For gene embedding comparison analyses, the maximum *PCC*_99.9_ cutoff generated among the permuted individual datasets was 0.2075. The set of 𝒢* genes for embedding comparison analysis were selected with cutoff ***PCC***^*^ >0.4, and the maximum feature embeddings per module was capped at ***e***_***max***_ = 25.

### Benchmarking against Hotspot

The Hotspot code was run as provided here https://hotspot.readthedocs.io/en/latest/Spatial_Tutorial.html, and the same preprocessed data was used as input for Smoothie. For runtime comparisons across different cell counts and gene counts, the same filtered count matrix was used for both Hotspot and Smoothie. Experiments were run on a high-performance computing system with an AMD EPYC 7763 64-core processor and 1000 GB of RAM.

## DATA AVAILABILITY

Supplementary data, including edge lists and node labels for all networks from this study, are available here https://doi.org/10.5281/zenodo.14933147.

## CODE AVAILABILITY

Smoothie is available here https://github.com/caholdener01/Smoothie, including easy to use example scripts.

## ACKNOWLEDGEMENTS

C.H. and I.D.V. would like to thank Dr. Sumanta Basu for helpful input on statistical testing of Smoothie and Dr. Yi Ren for helpful input on the ovary time course dataset. C.H. would also like to thank Nicole Eikmeier and Jerod Weinmann at Grinnell College for their network science and computer vision coursework that inspired the core idea of this paper. C.H. and I.D.V. would also like to thank all members of the De Vlaminck lab for helpful comments, discussion, and feedback. This work was supported by NIH R01AI176681, NIH R01AI176681, NIH R01AI189855 and CZI 2023-323354 to I.D.V.

## Author Contributions

C.H. and I.D.V. conceptualized the study. C.H. wrote the software. C.H. performed analyses with guidance from I.D.V.. C.H. and I.D.V. wrote the manuscript.

