## Supplemental Figures S1-8 for "Smoothie: Efficient Inference of Spatial Co-expression Networks from Denoised Spatial Transcriptomics Data"

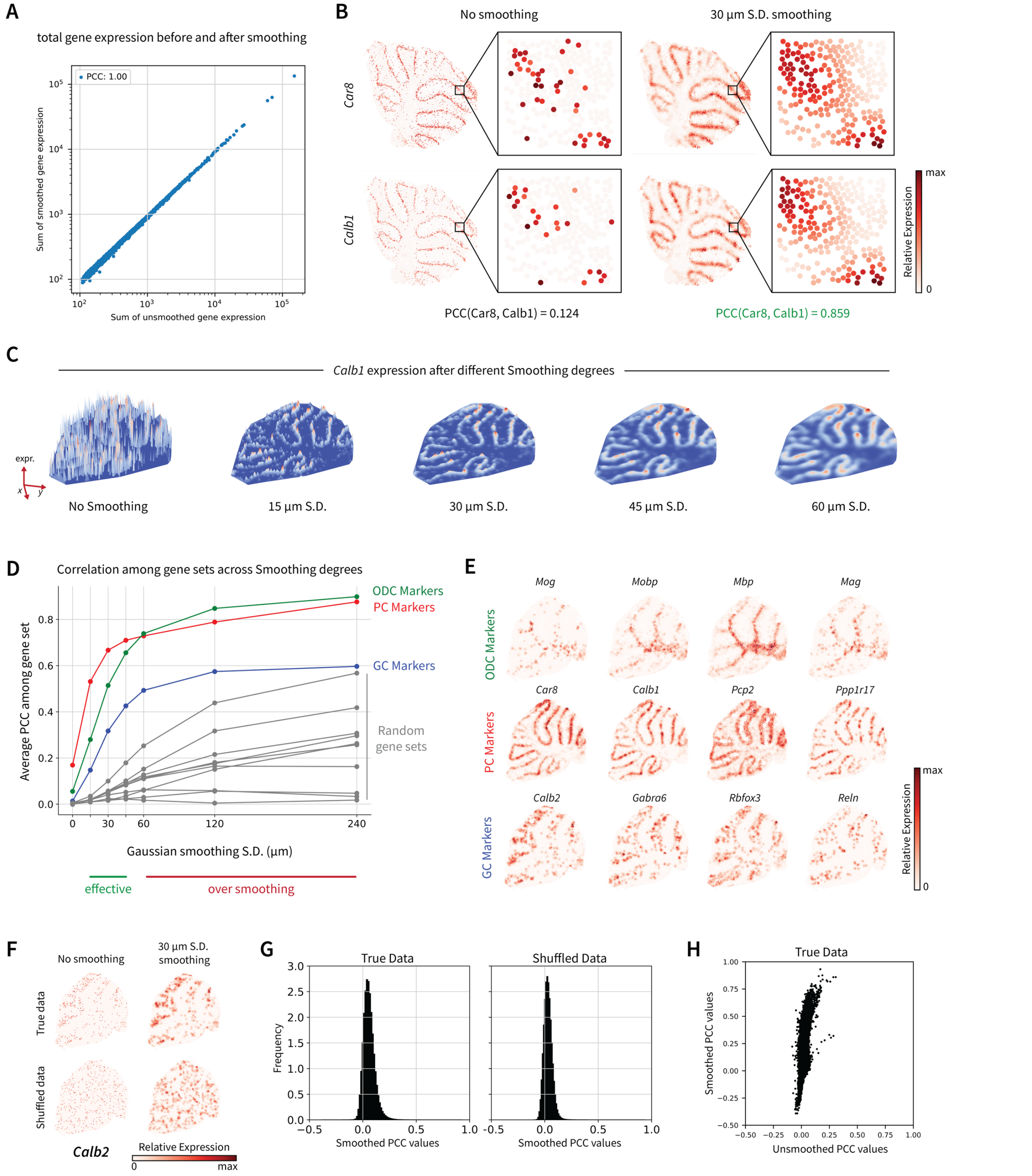


**Figure S1.** Implementation details for Gaussian smoothing and pairwise gene correlations. **A**. As a smoothing validation plot, gene expression sums before and after Gaussian smoothing are near perfectly correlated. **B**. Two known Purkinje cell marker genes *Car8* and *Calb1* have a low Pearson correlation coefficient (PCC) of 0.124 prior to smoothing due to fine-scale noise and RNA dropout. Post smoothing, the PCC value is raised to 0.859. **C**. 3D surface plots of the gene *Calb1* across different smoothing degrees. **D**. Average PCC values within various size-4 gene sets across different smoothing degrees. In the optimal smoothing range, PCC values among known marker gene sets are raised without over-inflating PCC values of random gene sets. **E**. Spatial plots of the gene sets from panel D, after 30 μm Gaussian standard deviation smoothing. **F**. Example of how gene *Calb2* looks after spatial permutation and then smoothing. **G**. The PCC distributions of all gene pairs in the smoothed true dataset versus the smoothed spatially shuffled dataset. **H**. After smoothing, the range of pairwise gene PCC values becomes much greater, allowing more accurate correlation measurements. (**F-H**. Gaussian s.d. = 30 μm).

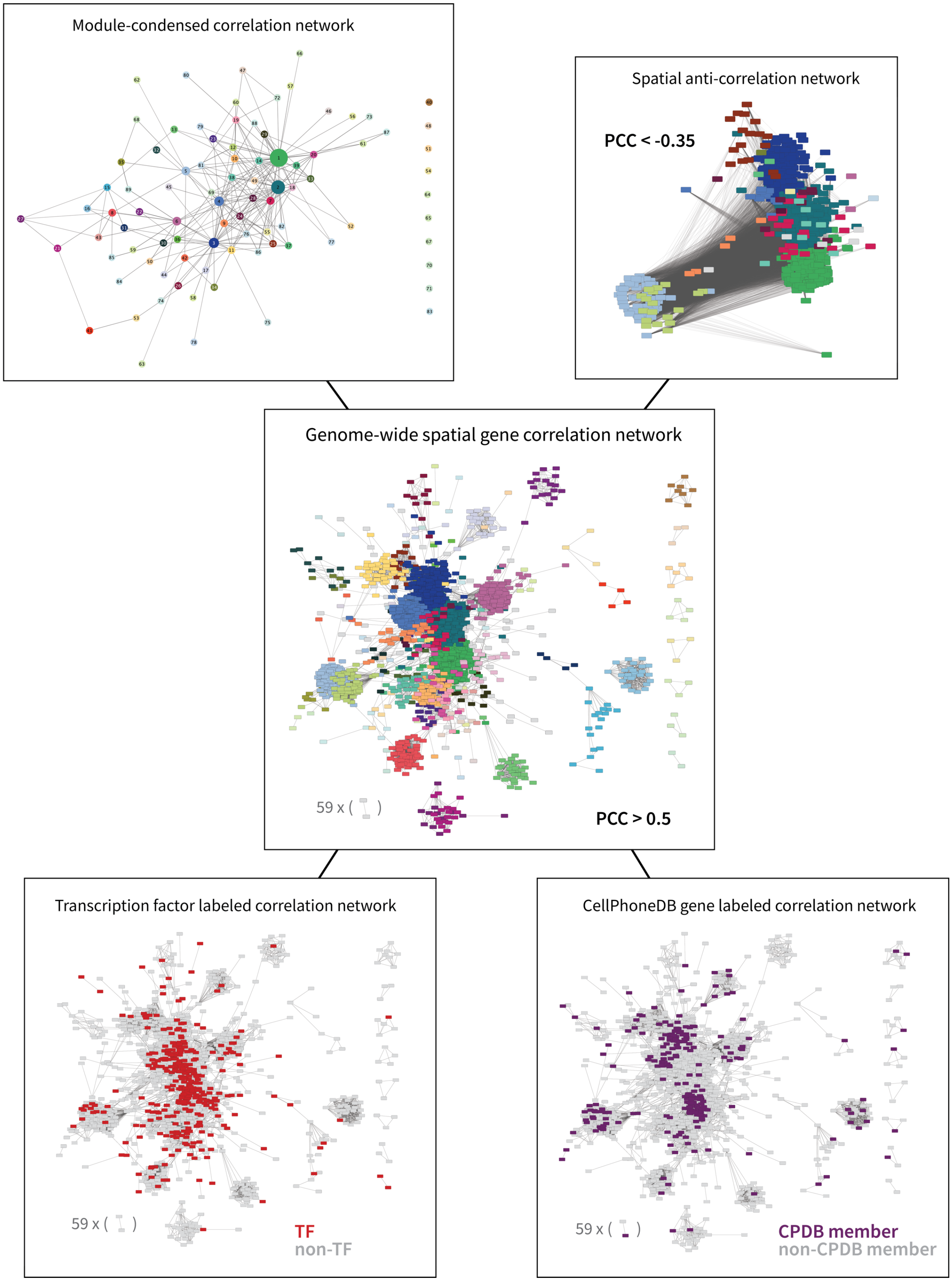


**Figure S2. Additional analyses from Smoothie's co-expression network output**. Module-condensed networks simplify the output to focus on how gene modules relate to each other. Anti-correlation networks reveal new information among genes and gene modules. Targeted investigations of transcription factors or cell-cell interaction genes (e.g. from CellPhoneDB) provide insight into gene regulation and cell communication occurring across the tissue.

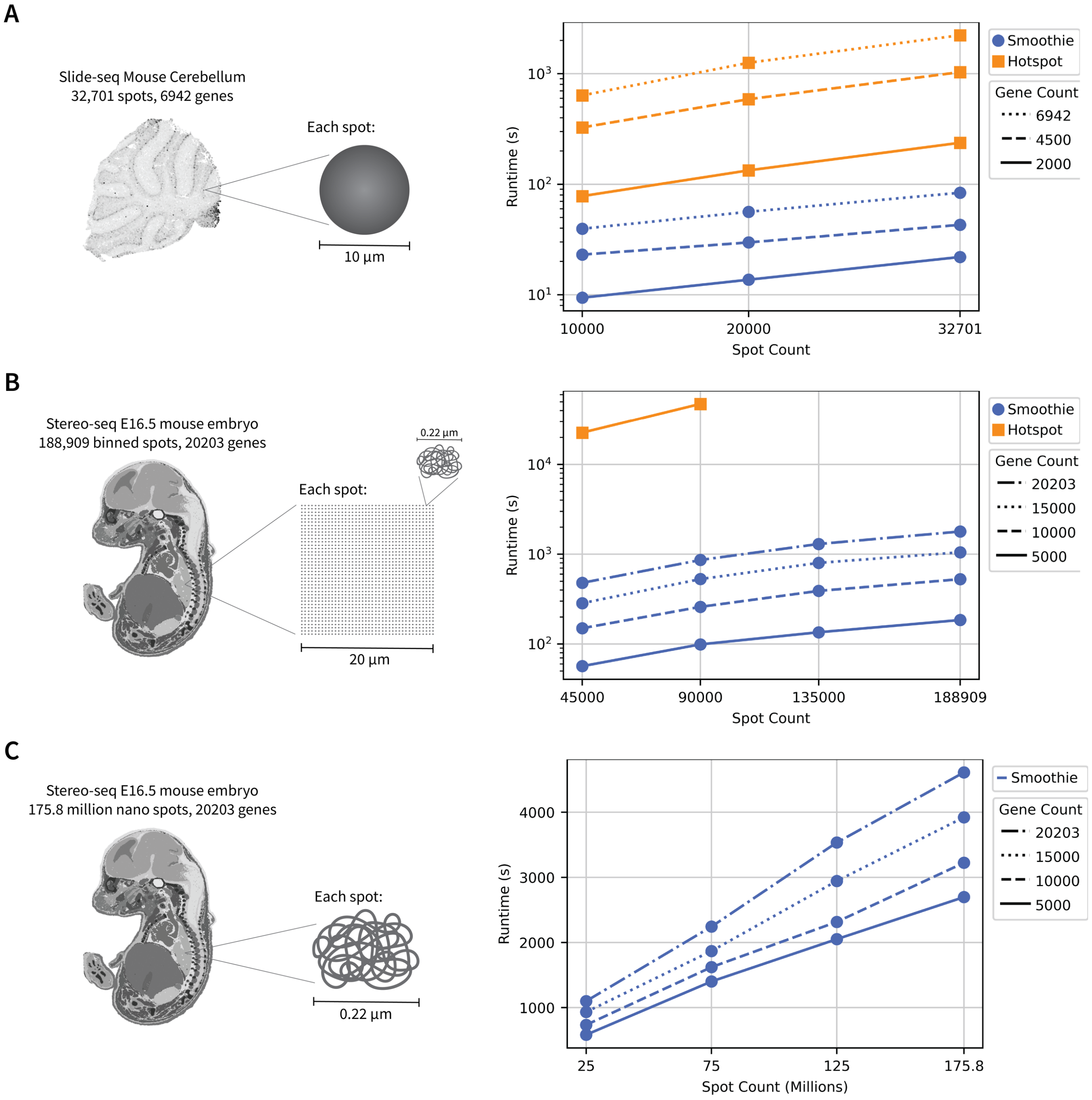


**Figure S3. Runtime benchmarking for Smoothie and Hotspot. A**. Runtimes for the two algorithms across different gene and spot counts in the 10-µm resolution Slide-seq cerebellum dataset, using 4 processes. **B.** Runtimes across gene and spot counts in the Stereo-seq mouse embryo dataset (20µm-binned), using 10 processes. Larger input experiments were excluded for Hotspot due to steep runtimes. **C.** Smoothie runtimes for grid-based smoothing on the millions of 220nm diameter Stereo-seq nano spots. Here, smoothing occurs directly over nano spots along a 20µm stride hexagonal point array. **A-C.** Across both algorithms, all experiments measure the time from the start of the algorithm up until outputting the pairwise gene correlation matrix, including all required statistical testing. For Smoothie, this entails smoothing and calculating the pairwise correlation matrix for both the true dataset and a spatially permuted version of the dataset (for significance testing). Hotspot parameters from the provided GitHub demo were used.

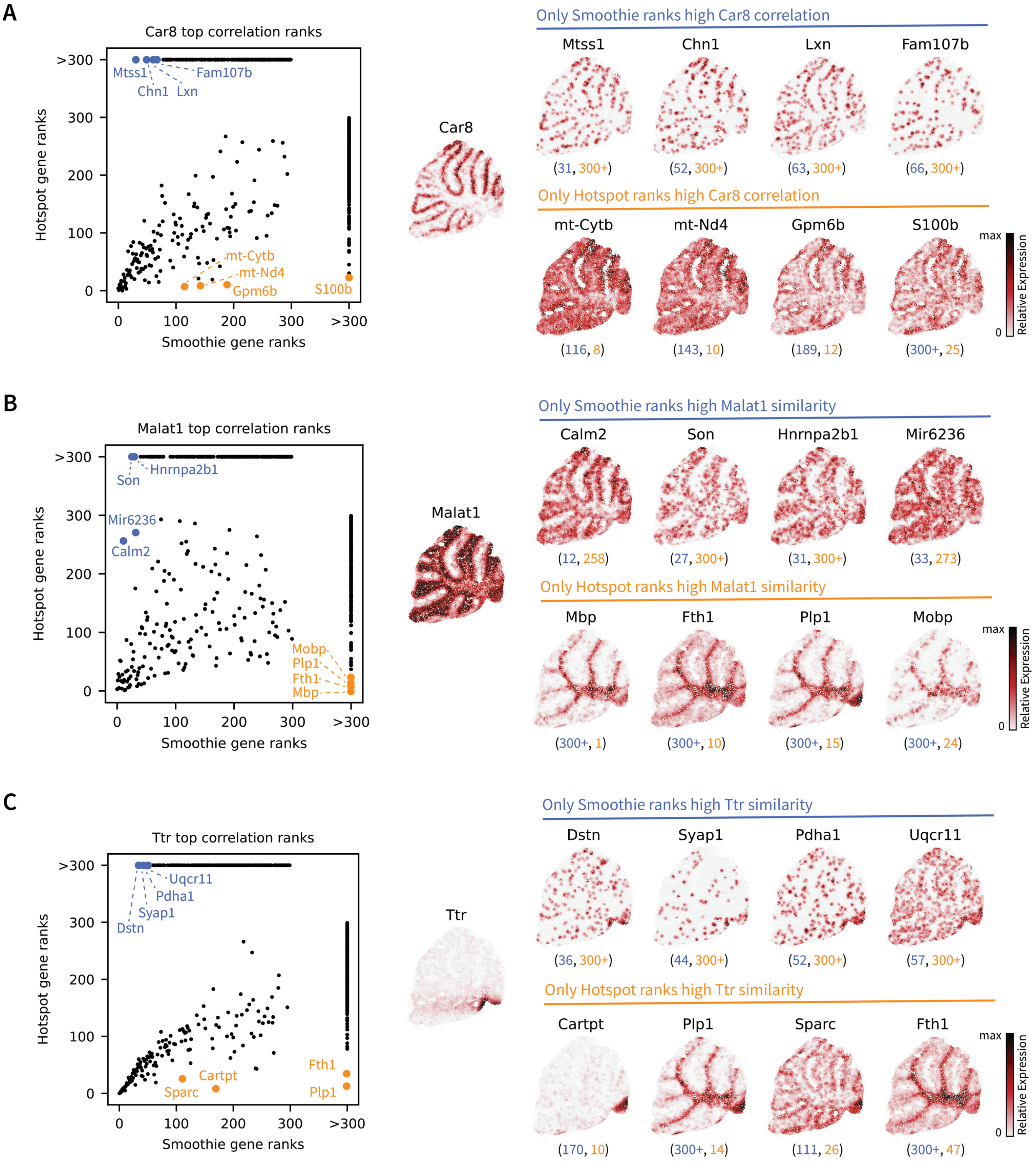


**Figure S4. Top gene correlation ranking comparison of Smoothie and Hotspot. A-C.** For genes *Car8*, *Malat1*, and *Ttr*, Smoothie and Hotspot largely agree on top ranking gene correlations, with some exceptions. Smoothie is better at detecting gene correlations with low-UMI, low-autocorrelation genes. Occasionally, Hotspot highly ranks gene patterns that have large differences from the target gene.

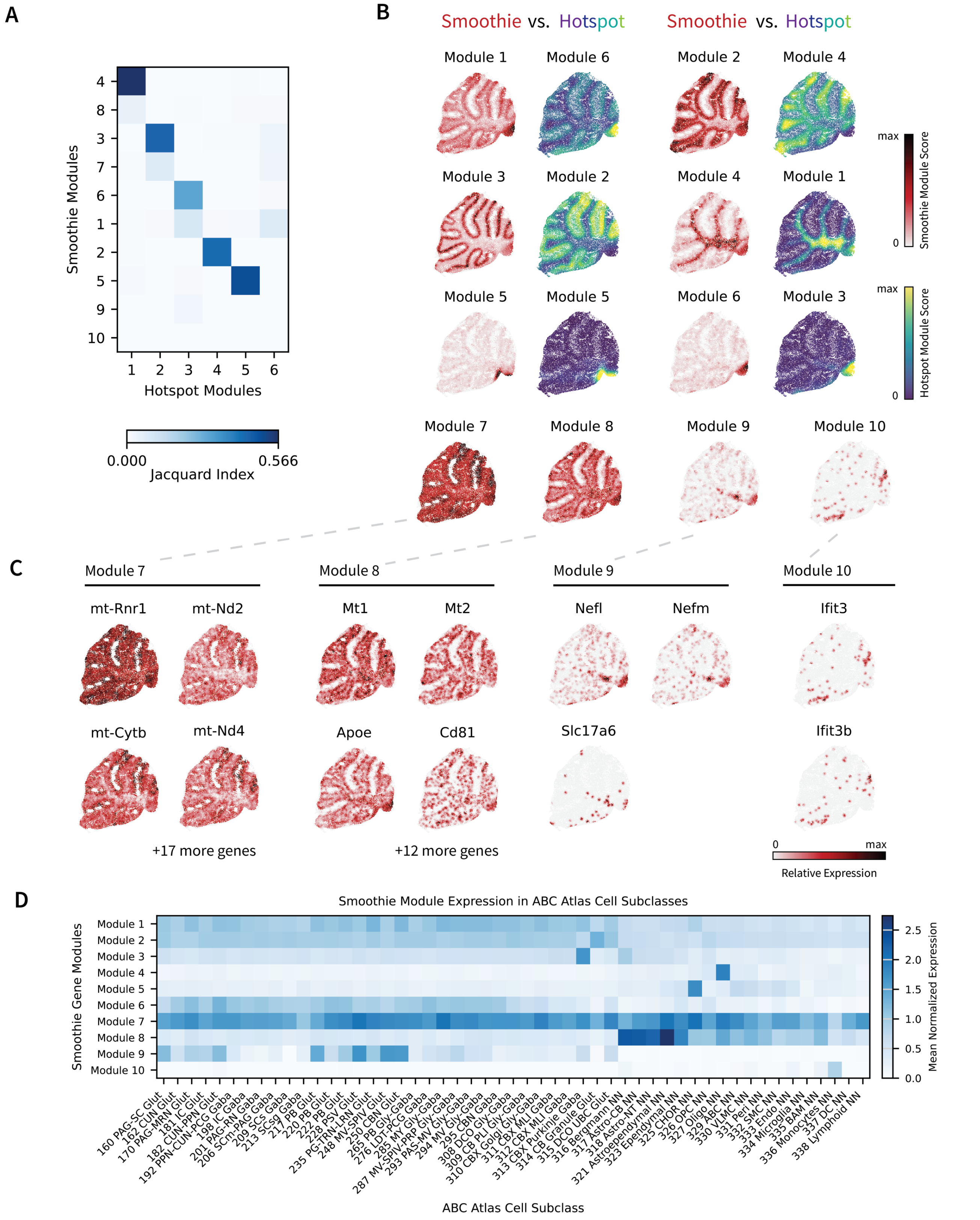


**Figure S5. Gene module comparison of Smoothie and Hotspot. A.** Jacquard index shows gene module overlap among Smoothie's 10 modules and Hotspot's 6 gene modules. **B.** Spatial plots of each method's gene modules show similarity among Smoothie's modules 1-6 and Hotspot's modules 1-6. **C.** Smoothie recovers 4 additional modules (7-10), which show distinct spatial patterns from modules 1-6 **D.** Plotting the Smoothie module's average expression in each cell subclass in the Allen Brain Cell (ABC) Atlas cerebellum region reveals meaningful connections of modules with cell type(s).

E16.5 Mouse Embryo Gene Pattern Atlas

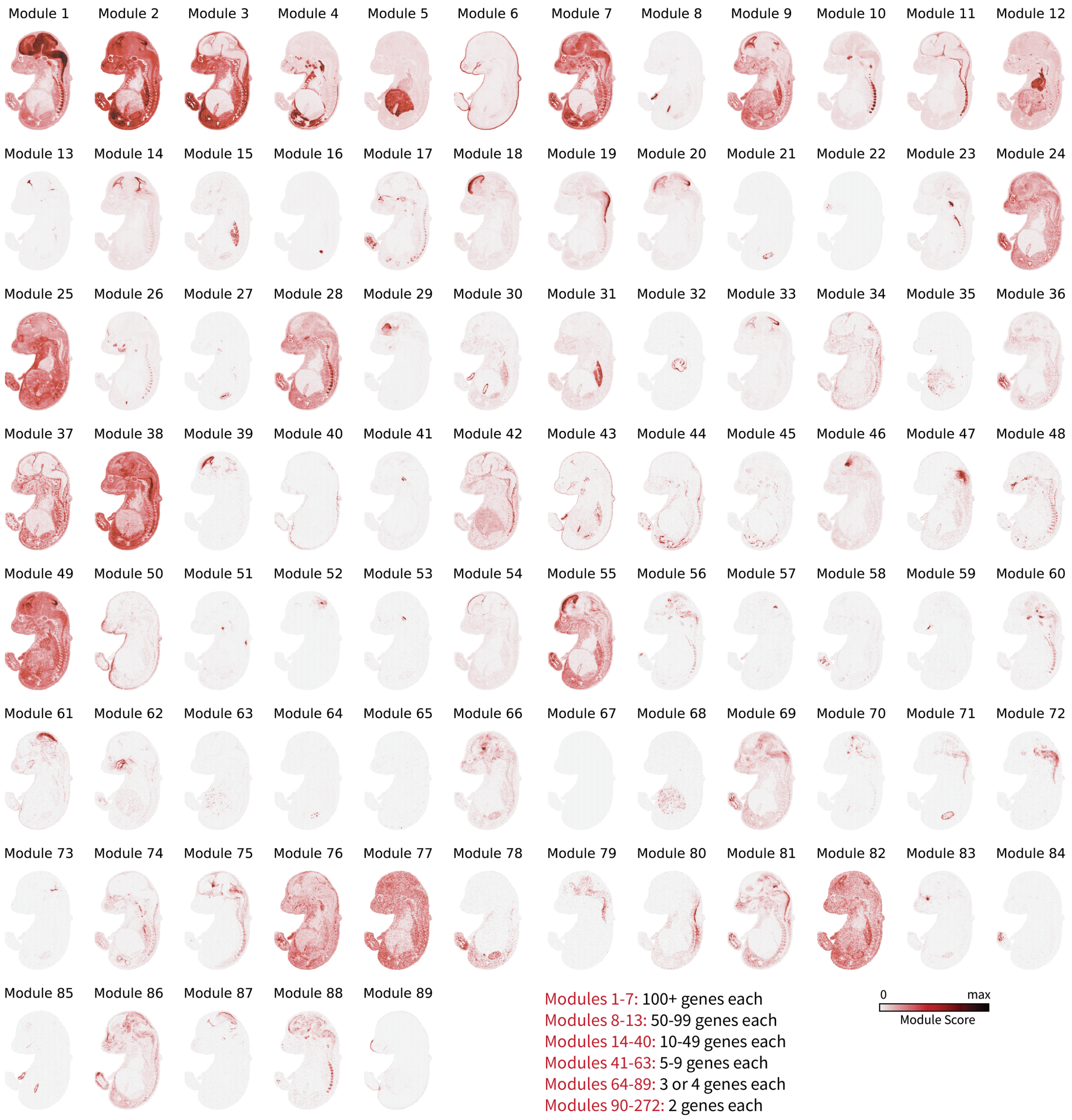


**Figure S6. Smoothie's spatial gene pattern atlas for the E16.5 Stereo-seq mouse embryo**. Lists of genes for each module are found in supplementary data, DataS4.

### Embryonic Day 13.5 to Day 16.5 Mouse Embryo Gene Pattern Atlas

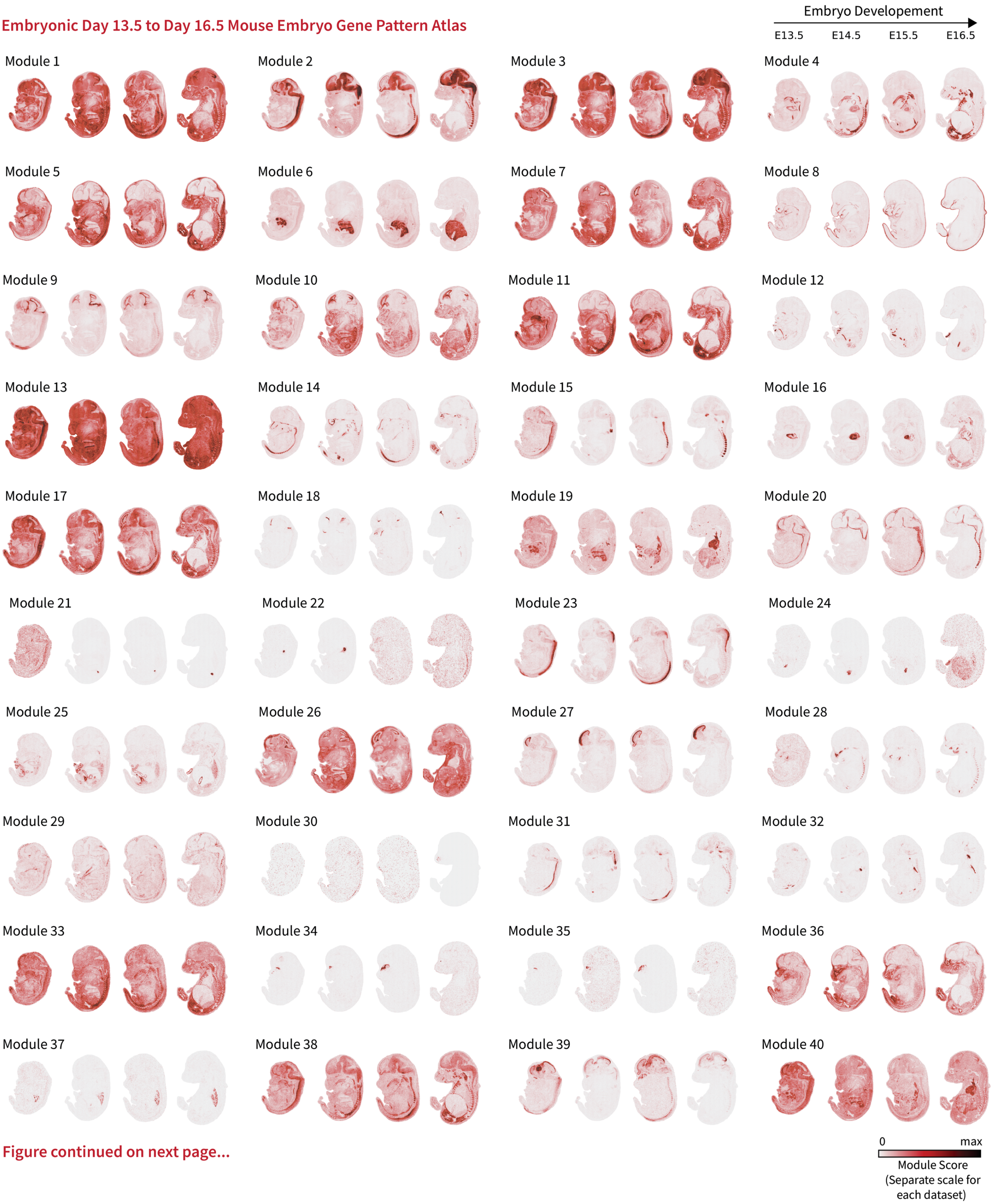


Figure continued on next page...

Embryonic Day 13.5 to Day 16.5 Mouse Embryo Gene Pattern Atlas (continued)

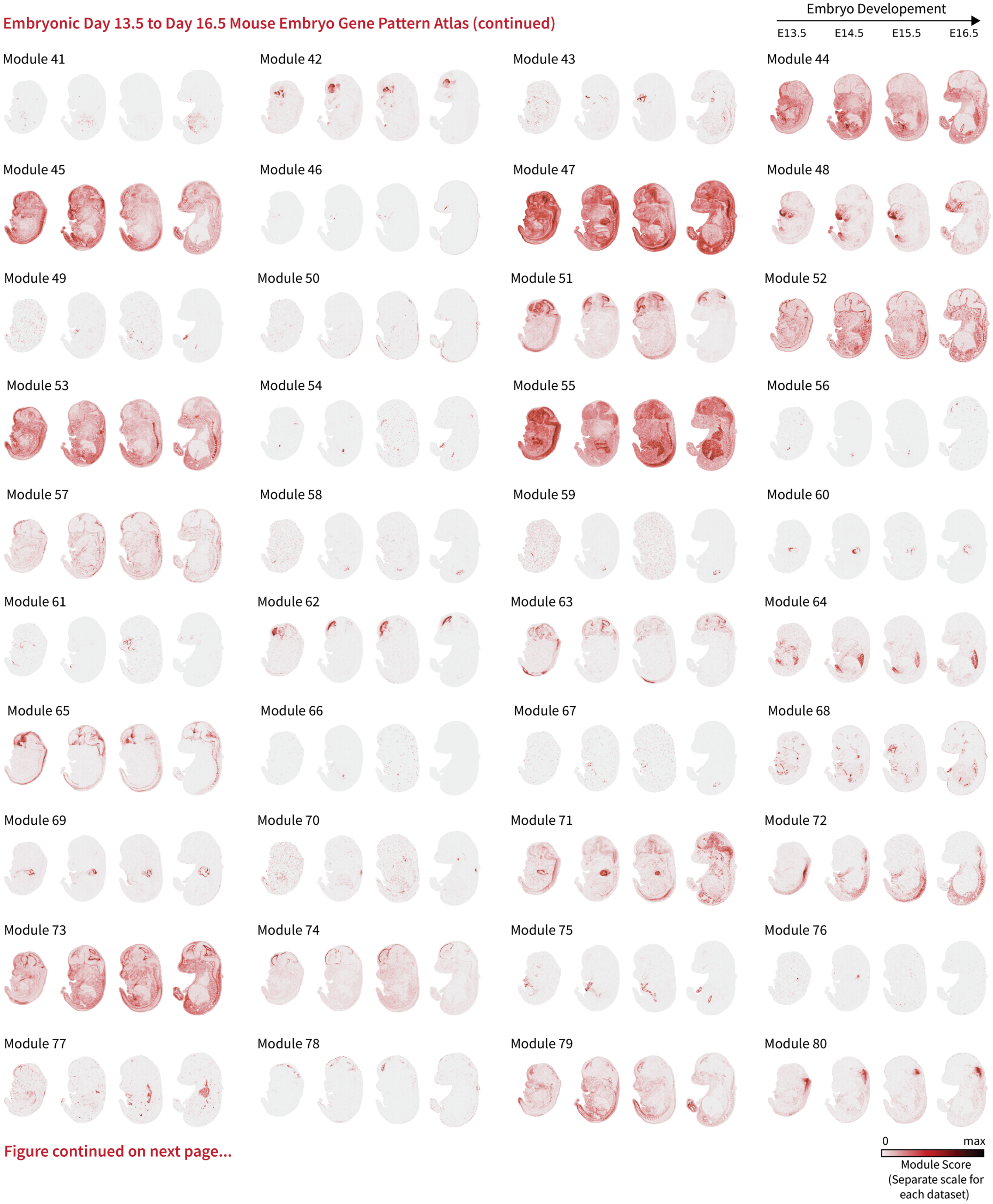


Figure continued on next page...

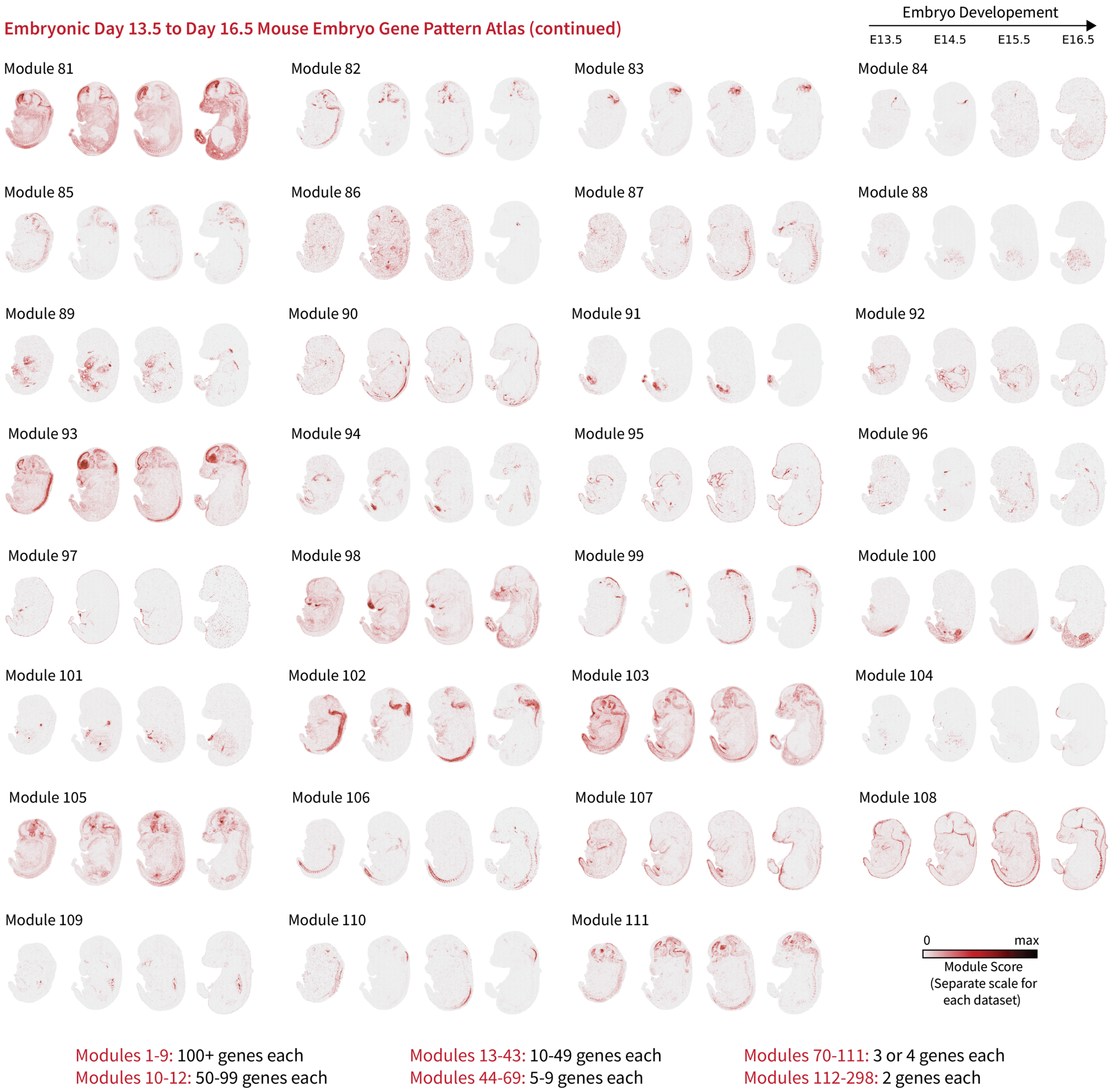


**Figure S7. Smoothie's spatial gene pattern atlas for the combined E13.5, E14.5, E15.5, and E16.5 mouse embryos dataset.** Lists of genes for each module are found in the supplementary data, DataS9.

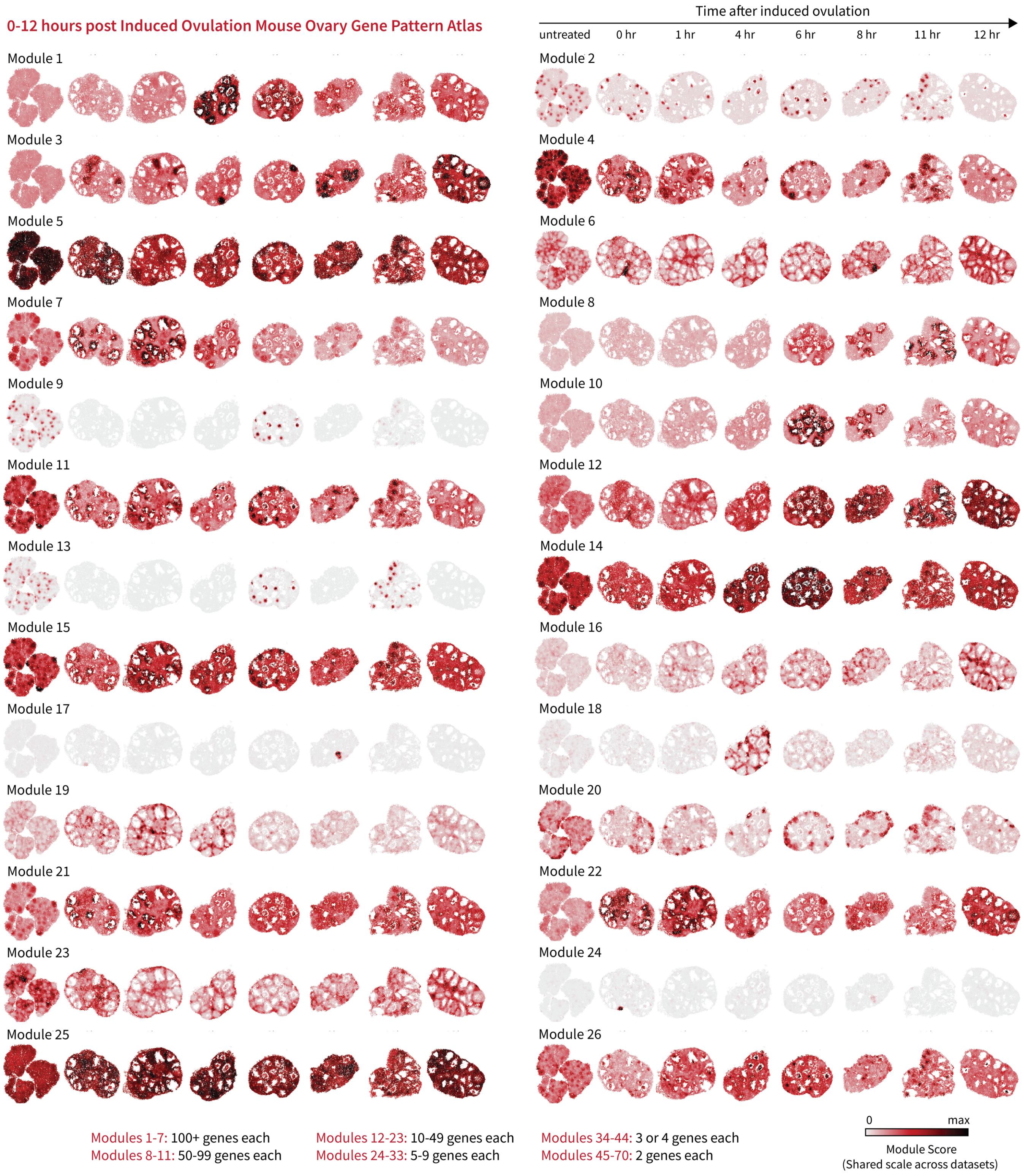


**Figure S8. Smoothie's spatial gene pattern atlas for the combined ovulating mouse ovary time course dataset.** Lists of genes for each module are found in the supplementary data, DataS12.
